# Highly Accurate Species Tree Inference from Orthologs and Paralogs through Scalable Quartet Amalgamation

**DOI:** 10.1101/2025.04.04.647228

**Authors:** Abdur Rafi, Ahmed Mahir Sultan Rumi, Sheikh Azizul Hakim, Md. Shamsuzzoha Bayzid

## Abstract

Species tree estimation from multi-copy gene family trees, including both paralogs and orthologs, is a challenging task due to gene tree discordance caused by biological processes such as incomplete lineage sorting (ILS) and gene duplication and loss (GDL). Summary methods, specially quartet-based species tree estimation methods, such as ASTRAL and quartet amalgamation-based methods in the Quartet MaxCut (QMC) and Quartet Fiduccia–Mattheyses (QFM) frameworks, have gained substantial popularity for their accuracy and statistical guarantee. However, most of the summary methods rely on single-copy gene trees and do not explicitly account for GDL, which limits their applicability to large genomic datasets containing multi-copy gene families. ASTRAL-Pro incorporates both orthology and paralogy for species tree inference under GDL by employing a refined quartet similarity measure based on the concept of speciation-driven quartets (SQs). In this study, we propose wQFM-GDL, the first successful initiative to model GDL in a quartet amalgamation-based summary method, using the SQ-based techniques effectively within the QFM framework. This required substantial algorithmic reengineering, including the development of efficient techniques for computing the initial bipartition in QFM and novel combinatorial methods for computing refined quartet scores directly from gene family trees. In addition, we introduce a locus-aware quartet weight normalization scheme for QFM framework under GDL, which significantly increased the accuracy of our method. We extensively evaluated our method, wQFM-GDL, on benchmark simulated and empirical datasets and compared it with leading methods, including ASTRAL-Pro3, SpeciesRax, FastMulRFS, and DupLoss-2. wQFM-GDL outperforms all other methods in 134 out of 156 model conditions considered in this study, with performance differences becoming more pronounced as dataset size increases. In particular, for larger datasets with 200 and 500 taxa, wQFM-GDL significantly outperforms all the competing methods across all 72 model conditions and achieves, on average, nearly a 25% reduction in reconstruction error compared with ASTRAL-Pro3. wQFM-GDL is freely available in open source form at https://github.com/abdur-rafi/wQFM-GDL.

## 1 Introduction

Species tree inference from genome-wide data is a central problem in evolutionary biology, but it is complicated by the fact that gene trees can differ from each other and from the underlying species tree. This phenomenon is known as gene tree discordance and may arise from several biological processes, including incomplete lineage sorting (ILS), gene duplication and loss (GDL), horizontal gene transfer, and hybridization (24). Among these processes, GDL plays a particularly important role in gene family evolution and frequently produces multi-copy gene family trees containing both orthologs and paralogs (32), which are widespread in phylogenomic datasets. Methods that require single-copy gene trees often restrict analyses to inferred single-copy orthologs or preprocess multi-copy gene families into single-copy subsets, thereby excluding a substantial fraction of the available genomic data from species tree inference (51).

Summary methods provide a scalable and accurate approach for species tree estimation in the presence of gene tree discordance. These methods combine the gene trees inferred individually from different loci or gene families to estimate a species tree. These methods are widely used due to their accuracy and statistical guarantee under particular reasons for gene tree discordance (2; 9; 17; 18; 19; 20; 21; 22; 25; 28; 31). Quartet-based summary methods form an especially important class of such approaches. A quartet is the evolutionary relationship induced by a tree on a set of four taxa, typically represented as a four-taxon unrooted binary tree. These methods address the Maximum Quartet Support Species Tree (MQSST) problem, which seeks to infer a species tree that maximizes the number of quartets induced by the gene trees that are consistent with the species tree.

Within this framework, methods can be broadly categorized by how they solve the MQSST problem. ASTRAL is the most widely used quartet-based summary method and uses dynamic programming to optimize quartet support within a constrained search space (28). In contrast, quartet amalgamation methods, such as QMC (Quartet Max-Cut) (2; 42), QFM (Quartet Fiduccia–Mattheyses) (26; 38), and SVDQuartet (9), use divide-and-conquer strategies to assemble quartets (with or without weights) induced by gene trees. However, traditional quartet amalgamation methods rely on explicit quartet generation and weighting, which can require Θ(*n*^4^) time for *n* taxa and becomes a major computational bottleneck for large phylogenomic datasets. Recent methods have addressed this scalability limitation by avoiding explicit quartet enumeration. TREE-QMC (15) and wQFM-TREE (36) compute the quantities required by their respective quartet-based optimization frameworks directly from gene trees. In particular, wQFM-TREE extends wQFM by retaining the same divide-and-conquer strategy, while replacing explicit quartet generation with direct combinatorial and graph-theoretic computations from gene trees. This substantially improves scalability while preserving the high accuracy of the wQFM framework. However, these methods were designed primarily for single-copy gene trees and therefore do not explicitly account for orthology and paralogy induced by gene duplication and loss.

A large family of summary methods considers GDL as a source of gene tree discordance. Many of them are reconciliation- or parsimony-based methods that seek a species tree minimizing the number of duplication and loss events or a related reconciliation cost. Examples include DupTree (45), iGTP (7), DupLoss-2 (33), DynaDup (3; 4; 5), and earlier dynamic-programming-based approaches (14). Other methods, such as PHYLDOG (6), Guenomu (10), MulRF (8), FastMulRFS (29), and SpeciesRax (30), address multi-copy gene families using different probabilistic, distance-based, or reconciliation-based formulations. Among quartet-based methods, ASTRAL-Pro (51) extended the ASTRAL framework to multi-copy gene family trees by defining a new quartet similarity measure that accounts for both orthology and paralogy. The key idea of ASTRAL-Pro is to consider only quartets that carry information about speciation events, defined as speciation-driven quartets (SQs) and ignore duplication-driven quartets having no speciation information. However, despite their effectiveness, no existing quartet amalgamation–based summary methods, such as wQFM and wQMC, explicitly model GDL and ASTRAL-Pro remains the only quartet-based summary method explicitly designed to account for GDL.

In this study, we introduce wQFM-GDL, an extension of the QFM framework that explicitly accounts for GDL and is the first initiative to model GDL in a quartet amalgamation method. We develop two variants of the method. The first, wQFM-GDL-T, extends wQFM-TREE to operate directly on rooted and tagged multi-copy gene family trees without explicitly enumerating quartets. To achieve this, we develop a new strategy for constructing initial bipartitions from multi-copy gene family trees and new combinatorial and graph-theoretic techniques for computing the speciation-driven quartets directly from gene family trees without explicit quartet enumeration. We further introduce a locus-aware normalization scheme that extends the quartet weight normalization used in the QFM framework to appropriately accommodate GDL. We also propose wQFM-GDL-Q where we enumerate the SQs from a given set of gene family trees and apply wQFM on this set of SQs. Notably, this set of SQs can be directly given as input to any quartet amalgamation methods (e.g., wQFM, wQMC). We found that, on small to moderate-sized datasets, where quartet enumeration is feasible, wQFM-GDL-Q is often more accurate than wQFM-GDL-T.

We evaluate wQFM-GDL on a broad collection of simulated and empirical datasets and compare it with leading methods for species tree inference from multi-copy gene family trees, including ASTRAL-Pro3, SpeciesRax, FastMulRFS, and DupLoss-2. Across model conditions varying duplication rates, loss rates, ILS levels, numbers of taxa, numbers of genes, and gene tree estimation error, wQFM-GDL achieves the best performance in most of the conditions considered in this study. In particular, on large simulated datasets comprising 200 and 500 taxa, wQFM-GDL significantly outperforms all competing methods in all 72 model conditions, achieving, on average, nearly a 25% reduction in reconstruction error relative to ASTRAL-Pro3. We also reanalyze three empirical datasets, Plants83 (46), Vertebrates188 (30), and Archaea364 (11), and find that wQFM-GDL successfully recovers all major relationships and does not violate any known consensus among scientists.

## 2 Materials and Methods

We first provide a brief overview of wQFM, wQFM-TREE and the duplication-aware quartet similarity measure used in ASTRAL-Pro. Then, we describe the key algorithmic components introduced in wQFM-GDL in detail.

### 2.1 Overview of wQFM and wQFM-TREE

wQFM (26) is a quartet amalgamation method that takes as input a set of weighted quartets induced by a given set of gene trees and infers a species tree by maximizing quartet consistency. The algorithm follows a divide-and-conquer strategy (Figure 1A): at each divide step, the current taxa set is partitioned into two disjoint subsets, defining two subproblems. For each divide step, wQFM starts from an initial bipartition on the current taxa set and iteratively refines it using a heuristic inspired by the Fiduccia–Mattheyses (FM) algorithm for hypergraph bipartitioning (13), with the objective of maximizing the difference between the number of satisfied and violated quartets. The procedure is applied recursively to the resulting subproblems until each subproblem contains at most three taxa, for which the solution is trivial. At each divide step, a dummy (artificial) taxon is introduced into each partition to represent the taxa in the complementary partition. During the conquer phase, the solutions to the subproblems (subproblem trees) are merged by connecting them through their corresponding dummy taxa, ultimately producing our final species tree.

**Figure 1:**
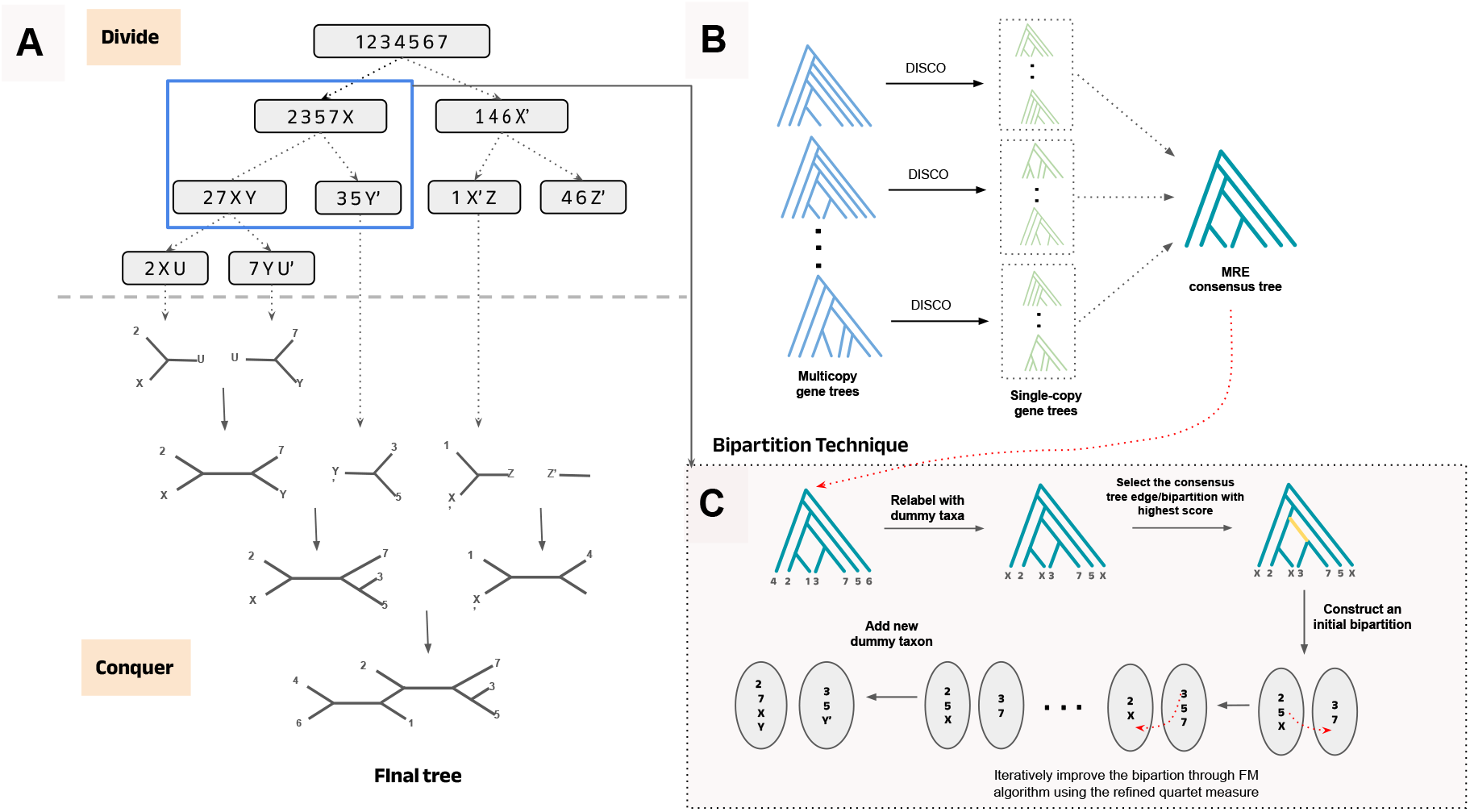
Overview of our proposed method. (A) The divide-and-conquer framework used in all QFM methods. (B) An MRE consensus tree is constructed from the multicopy gene trees after decomposing them into single-copy gene trees using DISCO which will be used to create an initial bipartition. (C) After generating the initial bipartition, it is iteratively improved using FM heuristic where the bipartitions are evaluated using the refined quartet measure addressing GDL.

Central to wQFM’s divide-and-conquer process is the identification of a suitable bipartition of the taxa set at each divide step. Each candidate bipartition is scored with respect to the input quartet set. A quartet *q* = ((*A, B*), (*C, D*)) is *satisfied* with respect to a bipartition (*P*_*a*_, *P*_*b*_) if taxa *A* and *B* lie in one partition and taxa *C* and *D* lie in the other, and is *violated* if taxa *A* and *C* (or *A* and *D*) lie in one partition while taxa *B* and *D* (or *B* and *C*) lie in the other. Otherwise, the quartet is *deferred*. The *score of a bipartition* is defined as the difference between the number of satisfied and violated quartets.

The main computational bottleneck of wQFM is the explicit enumeration of all quartets induced by the input gene trees. In earlier work, we developed wQFM-TREE (36) to address this limitation by enabling the wQFM framework to operate directly on gene trees without explicit quartet enumeration. This was achieved through novel algorithmic techniques that combine a gene tree consensus–based heuristic for constructing initial bipartitions at each divide step with combinatorial and graph-theoretic methods for computing the scores of candidate bipartitions directly from gene trees. We now revisit the normalization schemes used in TREE-QMC (15) and later adopted by wQFM-TREE, as they play an important role in the accuracy of the methods and are essential for understanding the new locus-aware normalization approach we propose in this study.

#### 2.1.1 Normalization in the QFM framework

Normalization is a critical component in the QMC and QFM frameworks because the divide-and-conquer process introduces artificial or dummy taxa into subproblems, which enables the subproblem trees to be merged in the conquer phase. Consequently, every non-root subproblem in the recursion contains one or more dummy taxa, whereas the input gene trees and the quartets induced contain only real taxa.

To compute satisfied and violated quartets with respect to a subproblem that contains dummy taxa, the leaves of each gene tree are appropriately labeled by dummy taxa. A dummy taxon represents all real taxa in its sister subproblem, and the corresponding leaves are relabeled using that dummy taxon. Thus, the resulting gene trees only contain the taxa present in the subproblem, including both real and dummy taxa. An important consideration is that, in this relabeled representation, multiple leaves of a gene tree may be relabeled by the same dummy taxon, which can inflate the contribution of quartets involving dummy taxa. However, from the perspective of the current subproblem, a dummy taxon should have the same influence as any single real taxon. To address this, quartet weights are normalized so that every subset of four taxa in a subproblem, including those containing dummy taxa, contributes a single vote toward the tally of satisfied or violated quartets.

Normalization was used in wQFM prior to TREE-QMC and wQFM-TREE but it was fairly simple because the full quartet set is available as input. Recently, TREE-QMC (15) introduced an effective normalization strategy which directly works on gene trees, and wQFM-TREE adopts a similar approach. Under this strategy, each taxon in a gene tree is assigned a weight and the weight of a quartet is defined as the product of the weights of its four taxa. Consequently, the total contribution of any quartet to the bipartition score reflects the effective representation of its constituent taxa in the current subproblem. The taxa that are absent from a subproblem but are represented through a dummy taxon receive appropriately reduced weights. This ensures that every subset of four taxa in a subproblem, including those containing dummy taxa, gets one vote per gene tree. The taxa are weighted non-uniformly depending on the subproblem decomposition in the recursion tree and taxa that are more relevant to a particular subproblem are assigned higher weights. The details of the normalization process with examples are presented in Supplementary Section 2.2. wQFM-GDL extends and improves the normalization scheme of wQFM-TREE to account for GDL (details in Section 2.3).

### 2.2 Overview of ASTRAL-Pro: Solving Maximum per-Locus Quartet-score Species Tree (MLQST)

The original ASTRAL tries to solve the NP-hard maximum quartet support species tree (MQSST) problem using an exact dynamic programming algorithm within a constrained version of the solution space. It tries to find a species tree that maximizes the number of quartets similar to the input gene trees. To account for orthology and paralogy, ASTRAL-Pro refines the quartet similarity measure in mainly two ways.

- Firstly, it only considers “speciation-driven quartets” (SQs) and excludes duplication quartets (DQs), which contain no information regarding speciation events.
- Secondly, it aggregates the speciation quartets that mandatorily share the same topology and separately do not provide any new information. These are counted as one unit, forming a quartet equivalence class.

Importantly, to classify quartets as SQs or DQs, the input trees must be rooted and tagged. Gene trees estimated from sequence data are neither rooted nor tagged. ASTRAL-Pro uses a maximum parsimony principle-based heuristic that tags each internal node in the gene trees as a speciation or duplication event. A quartet in a multicopy gene tree is classified as a speciation quartet (SQ) if, for every subset of three leaves, the corresponding most recent common ancestor (MRCA) is an internal node representing a speciation event (the rationale behind this definition is explained in detail in Supplementary Section 1). Now, the subtree of a binary tree restricted to a quartet exhibits two degree-3 nodes referred to as anchors and the MRCA/LCA of the two anchors of a quartet is called the anchor LCA. All the SQs on the same four species that share the same anchor LCA must also share the same topology (51). They are aggregated or counted as one unit to prevent double-counting. ASTRAL-Pro defines the per-locus quartet score of a species tree with respect to a gene family tree with tagged internal nodes to be the number of quartet equivalence classes of the gene trees agreeing with the species tree. Thus, it tries to solve the Maximum per-Locus Quartet-score Species Tree (MLQST) problem by reconstructing the tree having the maximum total per-locus quartet score with respect to the multicopy gene trees. Figure 2 illustrates examples of quartets in a gene tree that qualify as speciation quartets (SQs) and those that do not under this model. It also shows some examples of quartet equivalence classes. More details are presented in Supplementary Section 1.

**Figure 2:**
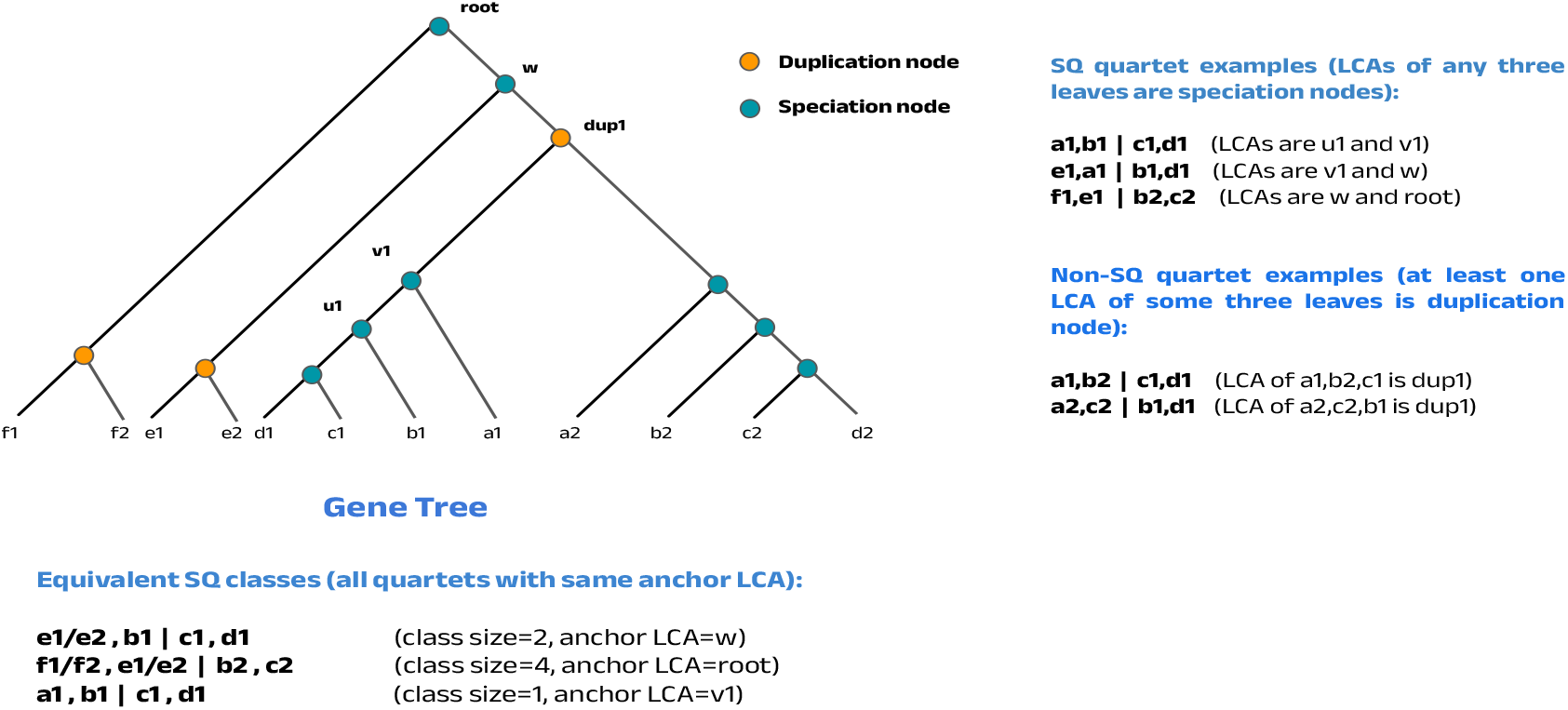
Examples of speciation-driven quartets (SQs), Non-SQs, and quartet equivalence classes.

### 2.3 wQFM-GDL-T: Extending wQFM-TREE for GDL

wQFM-GDL-T extends wQFM-TREE to accommodate GDL by solving the MLQST problem as in ASTRAL-Pro through three principal enhancements. First, we extend the consensus-tree-based heuristic to construct an appropriate initial bipartition of wQFM-TREE for multicopy gene trees. Second, we develop combinatorial techniques that enable efficient scoring of candidate bipartitions directly from multicopy gene trees without explicit quartet enumeration. Third, we enhance the normalization strategy of wQFM-TREE by proposing a locus-aware normalization scheme that explicitly accounts for GDL. An overview of the process is shown in Figure 1.

#### 2.3.1 Initial bipartition

At each divide step, we initiate the FM heuristic by generating an initial bipartition of the taxa set of the subproblem, which would then be refined iteratively, and the quality of this initial bipartition has a substantial impact on the quality of the final bipartition. In wQFM-TREE, an initial bipartition is obtained using a consensus-tree–based heuristic, in which an edge of a consensus tree constructed from gene trees is selected for initial bipartition. wQFM-TREE scores each bipartition according to its scoring scheme and selects the bipartition with the highest score. (36). However, for multicopy gene trees, a consensus tree is not directly defined. To address this limitation, we employ the method DISCO (49) to decompose multicopy gene trees into single-copy gene trees. We then construct a greedy consensus tree using the Majority Rule Extended (MRE) method implemented in the PAUP package (44), which supports incomplete gene trees as well. Then, we score each bipartition of the consensus tree using the updated scoring method addressing GDL as described later in this section and select the best one. The consensus tree is constructed only once in the beginning and used for all subproblems.

Importantly, an edge in the consensus tree cannot directly be used as a bipartition, since the taxa set of a subproblem may contain both real and dummy taxa (see Figure 1), whereas the consensus tree contains only real taxa. Therefore each edge/bipartition goes through a conversion process before being scored and considered as a candidate initial bipartition for that subproblem. The overall process is shown in Algorithm 1.

#### 2.3.2 Rooting and Tagging

Rooting and tagging is an essential step for quartet-based summary methods that address GDL. Quartet is the primary and natural unit of information in the unrooted setting, and earlier quartet-based methods worked directly on unrooted gene trees. However, in scenarios involving GDL, it is necessary to identify speciation and duplication events, which in turn requires rooting the gene trees. Gene trees inferred from sequence data, however, are typically neither rooted nor tagged. Therefore, ASTRAL-Pro uses a rooting and tagging procedure to root each gene tree and label each internal node as either a speciation or duplication node. For a given rooting, an internal node is considered a speciation node simply if the taxon sets of its left and right subtrees do not overlap; otherwise, it is considered a duplication node. The rooting is then chosen to minimize the number of duplications and losses.

However, this simple algorithm has some clear limitations. First, the idea that an internal node with no overlap between the taxa sets of its two child subtrees represents a speciation event can break down in the presence of ILS. In addition, duplication followed by a specially arranged pattern of losses can create scenarios where a node may represent a duplication event even though the taxa set of the subtrees do not overlap. Such scenarios are described as adversarial GDL (29). Moreover, assigning a single tag to each internal node may not always be appropriate, since some taxa in the left and right subtrees may be related through duplication while others may not. However, alternative strategies could be designed, which may include processes like rerooting and retagging gene trees using the species tree or probabilistic orthology inference are way more complex and future studies should explore those. Therefore, we use the rooting and tagging method of ASTRAL-Pro in wQFM-GDL.

##### Algorithm 1

Selection of Optimal Initial Bipartition in wQFM-GDL

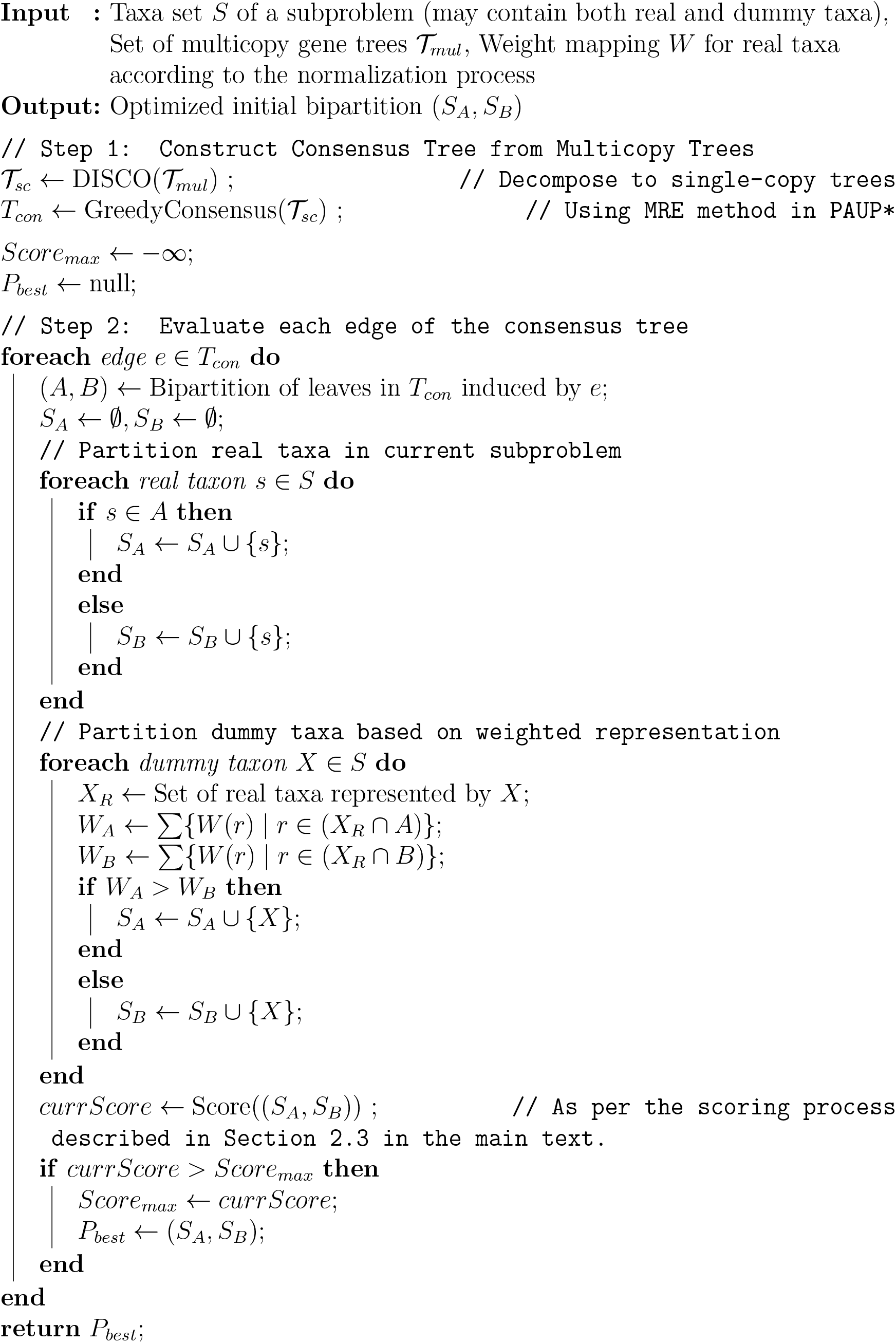

We believe that real biological datasets may contain complex GDL scenarios in which the existing tagging method may be less effective. Such scenarios may not be common in the traditional simulated datasets used to evaluate the methods. Future studies should investigate this issue in more detail.

#### 2.3.3 Scoring a Candidate Bipartition

After constructing the initial bipartition, we iteratively improve it. The taxa are transferred from one partition to the other iteratively following the FM heuristic to obtain progressively improved bipartitions. Each bipartition is scored to assess its quality. As described earlier, the score has been defined in the QFM framework as the difference between the total weight of quartets from the gene trees that the bipartition satisfies and violates.

The scoring mechanism in wQFM-GDL differs substantially from wQFM-TREE, as it must account for the complexities introduced by GDL. In the original wQFM-TREE, all quartets from single-copy gene trees are considered when computing the score. In contrast, for multicopy gene family trees, following the quartet similarity measure of ASTRAL-Pro, we restrict the calculation of satisfied and violated quartets only to speciation-driven quartets (SQs) and treat each quartet equivalence class as a single unit to avoid double-counting. To integrate these duplication-aware measures while preserving the scalability of the original wQFM-TREE, we had to introduce substantial modifications to the underlying combinatorial and graph-theoretic techniques of wQFM-TREE to enable the exclusion of all the duplication-driven quartets and the aggregation of equivalent quartets within the scoring framework. The extended terminology and the updated scoring procedure are described below.

Let G be a set of rooted and tagged multi-copy gene family trees with taxa set X. Multiple leaves corresponding to different copies from the same taxon carry the same label. We root and tag the gene trees using the algorithm of ASTRAL-Pro. *ab*|*cd* denotes a quartet induced by four selected leaves labeled *a, b, c, d*. Let (*A, B*) be a candidate bipartition of the taxa set *S* in a divide step. We define *R*_*A*_ and *D*_*A*_ as the sets of real and dummy taxa in *A*, respectively. We define *F*_*A*_ as the set of all real taxa that are either in *R*_*A*_ or represented by a dummy taxon *X* ∈ *D*_*A*_, that is, 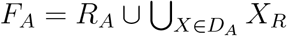. We similarly define *R*_*B*_, *D*_*B*_, and *F*_*B*_.

Now, a quartet *ab*|*cd* in a rooted and tagged gene family tree is an SQ only if *a, b, c, d* represent four distinct taxa, i.e., four distinct labels from X and the MRCAs of all triplets of the selected leaves labeled *a, b, c, d* are speciation nodes in its respective gene tree. For scoring a candidate bipartition in the current subproblem, we further consider only those SQs for which no two of the four taxon labels are represented by the same dummy taxon. This ensures that a dummy taxon, which represents a single taxonomic unit in a particular subproblem, does not contribute more than once to the same quartet. We categorize the considered SQs using the same traditional approach as in the QFM framework.

1. **Satisfied SQ:** SQ *ab*|*cd* is satisfied if *a, b* ⊆ *F*_*A*_, *c, d* ⊆ *F*_*B*_ or vice versa.
2. **Violated SQ:** SQ *ab*|*cd* is violated if the bipartition separates the four taxon labels according to one of the two alternative quartet topologies, *ac*|*bd* or *ad*|*bc*. Equivalently, either *a, c* and *b, d* lie on opposite sides of the bipartition, or *a, d* and *b, c* lie on opposite sides of the bipartition.
3. **Deferred SQ:** Any other SQ.

Now, our algorithm assigns weight to each real taxon in X according to the normalization process (Supplementary Section 2.2). We consider a quartet as composed of two unordered pairs *a, b* and *c, d*. The weight of a pair, *w*(*a, b*) = *w*(*a*) · *w*(*b*) and weight of a quartet *ab*|*cd* is defined as *w*(*ab*|*cd*) = *w*(*a, b*) · *w*(*c, d*) = *w*(*a*) · *w*(*b*) · *w*(*c*) · *w*(*d*). The weight of a set *Q* of quartets is defined as *w*(*Q*) = ∑_*q*∈*Q*_ *w*(*q*). The weights vary across subproblems according to the normalization process.

Let 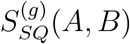 and 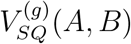 denote the sets of satisfied and violated SQ equivalence-class representatives, respectively, with respect to the candidate bipartition (*A, B*), each containing only one representative from each equivalence class in a gene tree *g* ∈ G. The score of a candidate bipartition (*A, B*) with respect to G is defined as:

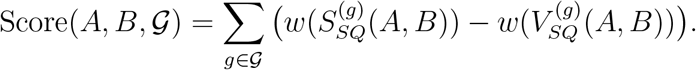

In this setting, computing the quantities 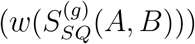 and 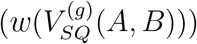 are more complex and different than wQFM-TREE. The score must be restricted to SQ equivalence-class representatives: which quartets qualify as SQs depends on the speciation/duplication tagging of the gene family tree, while equivalence among SQs is determined by the four taxon labels and the anchor LCA associated with each SQ. Therefore, computing these quantities directly from gene family trees requires new combinatorial and graph-theoretic techniques that exclude duplication-driven quartets and aggregate equivalent SQs without explicitly enumerating all quartets. We describe this computation below.

##### Computing 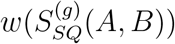 for Multi-copy Trees

Our key observation is that the total weight of all satisfied SQs sharing the same anchor LCA (Least Common Ancestor (LCA) of the two anchors of a quartet) can be computed efficiently and independently, and then aggregated for all anchor LCAs. The anchor LCA of a quartet is the least common ancestor of its two anchors. Since a quartet can be an SQ only when the MRCAs of all triplets of its four selected leaves are speciation nodes, a duplication node cannot serve as the anchor LCA of an SQ. Therefore, we only iterate over the nodes tagged as speciation in the gene family tree.

Rooted quartets exhibit two distinct forms: balanced and unbalanced, as shown in Figure 3A. We observe that, for each particular anchor LCA, we can efficiently calculate the balanced and unbalanced satisfied SQs separately. Let *u* be an internal node tagged as speciation with child branches *i* and *j*. For *Y* ∈ {*A, B*} and *l* ∈ {*i, j*}, we define 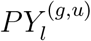 as the set of unique unordered pairs *a, b* of distinct taxa such that *a, b* ∈ *F*_*Y*_, selected leaves labeled *a* and *b* descend from branch *l* of *u*, and the two taxa are not represented by the same dummy taxon in the current subproblem.

**Figure 3:**
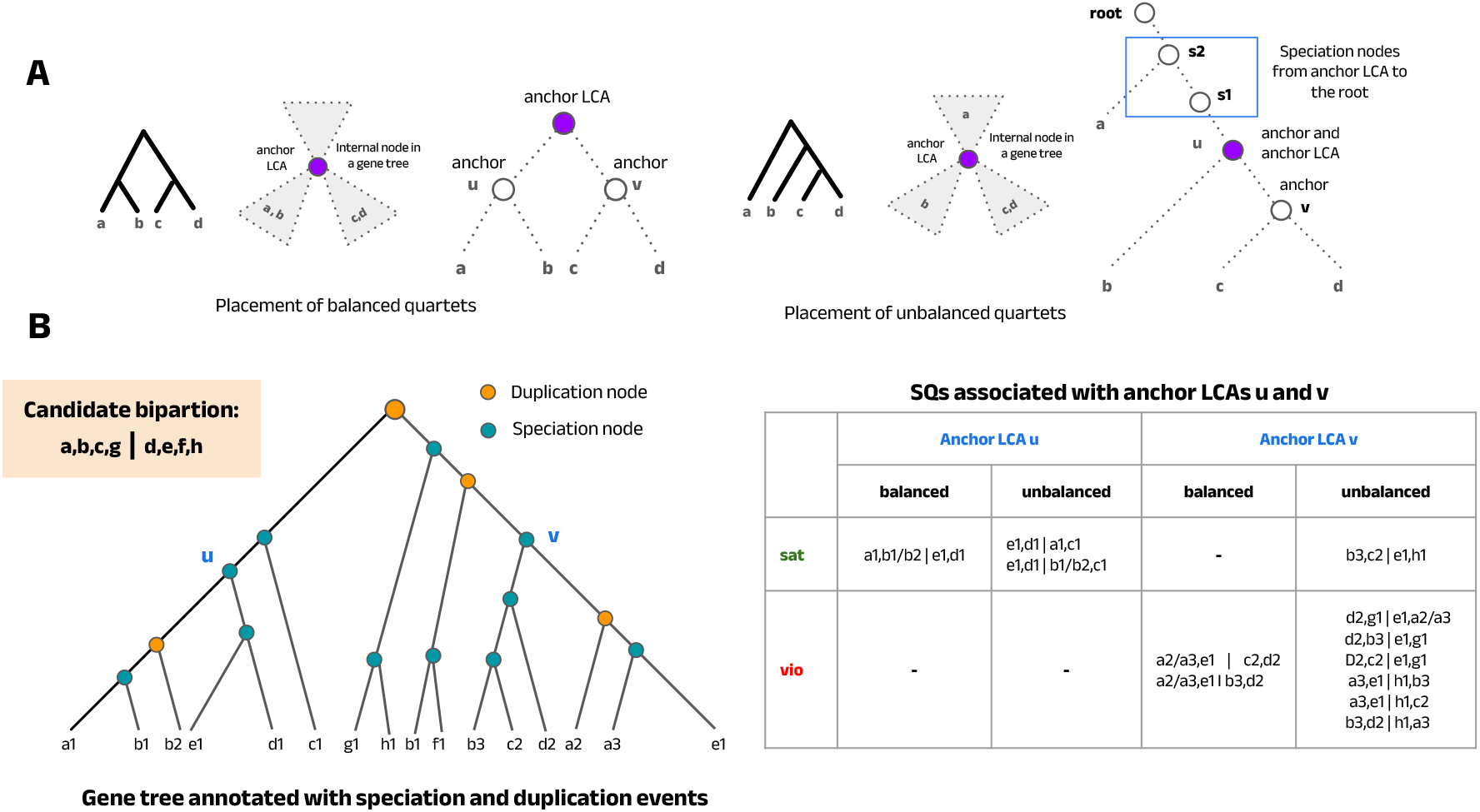
Scoring a candidate bipartition. (A) Placement of a balanced and unbalanced quartet in a gene tree with respect to their anchor LCA. (B) SQs of different categories, illustrated using anchor LCAs *u* and *v* for an example bipartition *a, b, c, g*|*d, e, f, h*. Each speciation node may act as an anchor LCA for some SQs and for scoring a bipartition, we compute the total weight of satisfied and violated SQs associated with each anchor LCA independently, treating balanced and unbalanced quartets separately and then sum the results.

Importantly, multiple identical pairs may arise within both the left and right subtrees, but we count only one representative instance. When pair representatives are combined to form quartets associated with anchor LCA *u*, different choices of duplicated leaves may involve the same four taxa and share the same anchor LCA. Such quartets belong to the same SQ equivalence class and are therefore counted only once. This ensures that each SQ equivalence class contributes a single representative to the score, consistent with the ASTRAL-Pro quartet similarity measure. This is a key distinction from the original wQFM-TREE calculation, where all quartets induced by single-copy gene trees are considered.

##### Balanced Structure

A rooted quartet is balanced if the root of the quartet separates its four leaves into two pairs. A balanced SQ with anchor LCA *u* has one anchor in branch *i* and the other in branch *j*, and each anchor subtends one of these two pairs in the restricted quartet (Figure 3A). The SQ is satisfied if the pair from one branch belongs to *F*_*A*_ and the pair from the other branch belongs to *F*_*B*_. Let 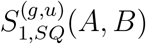 denote the set of balanced satisfied SQ equivalence-class representatives with anchor LCA *u*. Thus, we compute

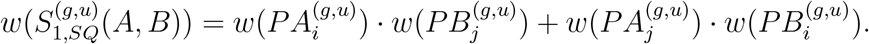

##### Unbalanced Structure

A rooted quartet is unbalanced if the root of the quartet separates one leaf from the other three, creating a ladder-like structure. For an unbalanced SQ with anchor LCA *u*, one anchor lies inside one of the child branches of *u*, while the other anchor is *u* itself. Suppose the lower anchor lies in branch *i*, without loss of generality. In this case, two taxa are selected from branch *i*, one taxon is selected from the sibling branch *j*, and the fourth taxon is selected from the parent side of *u* or outside the subtree rooted at *u* (Figure 3A).

However, the fourth taxon cannot be chosen arbitrarily from the parent side. To preserve the SQ condition, it must be descendant to a branch associated with a speciation node on the path from *u* to the root (Figure 3A); otherwise, the MRCA of some triplet of the four selected leaves may be a duplication node. Therefore, the fourth taxon reached from *u* via its parent edge *k* is only from this eligible region of the parent side. Thus, for a fixed anchor LCA *u*, an unbalanced SQ contains either a pair from branch *i* and a pair split between branch *j* and branch *k*, or a pair from branch *j* and a pair split between branch *i* and branch *k*.

We define 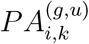 as the set of unique unordered pairs *a, b* of distinct taxa in *F*_*A*_. In each pair, one taxon descends from branch *i*, and the other belongs to the eligible parent-side region *k*. We exclude pairs in which both taxa are represented by the same dummy taxon in the current subproblem. The set 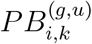 is defined analogously for taxa in *F*_*B*_. The sets 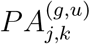 and 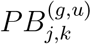 are defined similarly.

Let 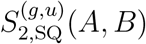 denote the set of unbalanced satisfied SQ equivalence-class representatives with anchor LCA *u*. Then the total weight of these representatives is computed as

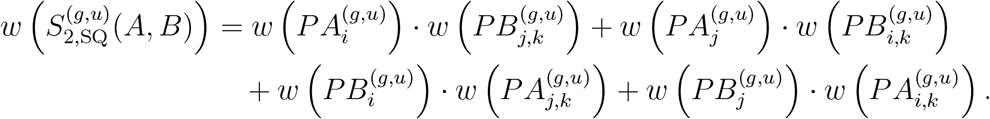

Since balanced and unbalanced rooted quartet structures are disjoint, the total weight of satisfied SQ equivalence-class representatives associated with anchor LCA *u* is

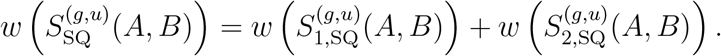

We then aggregate over all speciation nodes of *g*:

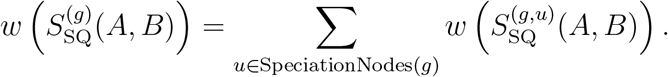

An example of this process is shown in Figure 3.

Here, we observe that duplication-derived quartets in the gene trees are effectively excluded, as the LCA of at least one triplet among the four taxa of these quartets corresponds to a duplication node and therefore does not satisfy any of the criteria mentioned for getting considered in both balanced and unbalanced quartet calculations.

##### Computing 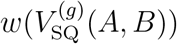 for Multi-copy Trees

The calculation of 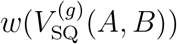 follows the same anchor-LCA-based decomposition used for satisfied SQs. For each speciation node *u*, we compute the total weight of violated SQ equivalence-class representatives associated with anchor LCA *u* by considering balanced and unbalanced structures separately, and then aggregate the contributions over all such nodes.

For balanced rooted quartets, the gene-tree topology places one pair of taxa in branch *i* and one pair of taxa in branch *j*. If the quartet is violated by the candidate bipartition (*A, B*), in each branch, one of the taxa contained in *F*_*A*_ and the other contained in *F*_*B*_. Thus, the quartet contains two taxa from *F*_*A*_, one from each of its child subtrees, and similarly two taxa from *F*_*B*_, one from each of its child subtrees. Let 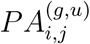 denote the set of unique unordered pairs *a, b* of distinct taxa such that *a, b* ∈ *F*_*A*_, one taxon descends from branch *i*, the other descends from branch *j*, and the two taxa are not represented by the same dummy taxon in the current subproblem. The set 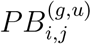 is defined analogously for taxa in *F*_*B*_.

Let 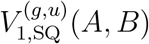 denote the set of balanced violated SQ equivalence-class representatives with anchor LCA *u*. Then

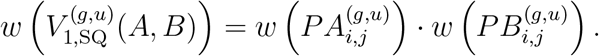

Similarly, we can compute the weight of violated unbalanced quartets. Let 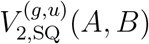 denote the set of unbalanced violated SQ equivalence-class representatives with anchor LCA *u*.

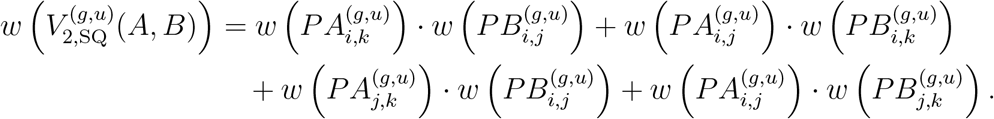

Since balanced and unbalanced rooted quartet structures are disjoint, the total weight of violated SQ equivalence-class representatives associated with anchor LCA *u* is

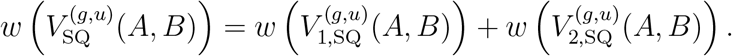

We then aggregate over all speciation nodes of *g*:

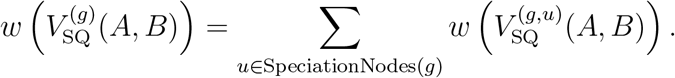

###### Theorem 1

(Correctness of anchor-LCA-based SQ scoring). *For any rooted and tagged multi-copy gene family tree g and candidate bipartition* 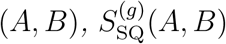 *and* 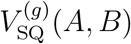 *include only the satisfied and violated SQs, respectively and exclude DQs. Moreover, they contain only one quartet from each equivalent class*.

*Proof*. We prove the result using three claims.

*Claim 1:* 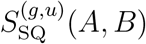 *contains precisely the satisfied SQ equivalence classes whose anchor LCA is u*.

Consider any SQ equivalence class representative induced by *g* with anchor LCA *u*. By definition, it contains four distinct taxa, and the MRCA of every triplet of the corresponding leaves is a speciation node. Therefore, *u* must be a speciation node of *g*. Thus, all of the quartets in the class are considered by the computation of balanced or unbalanced quartets followed for 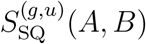 and 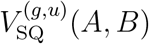. Moreover, it cannot be included by any other 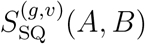 component, where *v*≠ *u*, which proves our claim.

*Claim 2:* 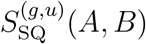 *and* 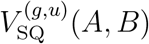 *include only one quartet of every satisfied or violated SQ equivalence class with anchor LCA u*.

First, all quartets within an SQ equivalence class have the same weight. Since they involve the same four species, if one quartet in the class is satisfied (or violated) by a bipartition, then all other quartets in the class are also satisfied (or violated), respectively. Furthermore, all quartets in the class are either balanced or unbalanced simultaneously. Therefore, from the perspective of a fixed anchor LCA *u*, the quartets within an equivalence class are indistinguishable, and exactly one representative is counted through either 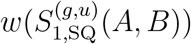 or 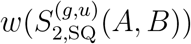. Now, for an equivalence class with balanced quartets, they contain one pair of taxa from the left subtree of *u* and the other from the right subtree of *u*. In the unbalanced case, for all equivalent quartets, one pair is from one of the subtrees, while the remaining two taxa come from the other subtree and a selected region of the parent branch. Here, when calculating the total weights, the pair sets retain only one representative for each unordered taxon pair. Similarly, for the unbalanced quartets, where a single taxon is selected from a subtree and parent branch, duplicates are counted only once. Different choices of duplicate taxon pairs and singleton taxa within the subtrees induce the same set of four taxa and, consequently, quartets belonging to the same SQ equivalence class. Since identical unordered taxon pairs and singleton taxa are counted only once, 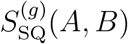 and 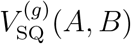 contain exactly one representative from each satisfied or violated SQ equivalence class with anchor LCA *u*.

*Claim 3: No non-SQ quartet is counted*.

The calculation only considers speciation nodes as possible anchor LCAs. For unbalanced quartets, the fourth taxon is further restricted to a selected region of parent branch *k*, ensuring that the MRCA of every triplet remains a speciation node. Specifically, it must come from a branch associated with a speciation node on the path from the anchor LCA to the root. Therefore, any quartet for which some triplet has a duplication-node MRCA is excluded. Thus, every quartet representative counted by the formulas satisfies the SQ condition and is valid for the current subproblem.

Thus, no non-SQ quartet is considered, and among the SQs, only one representative from each equivalence class is counted and exactly once. This completes our proof.

Now, computing the pair weights 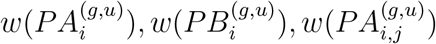, and the analogous terms without enumerating the quartets requires additional combinatorial techniques and efficient graph traversal algorithms, which we describe next.

##### Computing weights of taxon pairs efficiently

The components 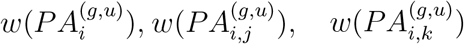, and the analogous terms represent weights of taxon pairs, which are combined to get the quartet weights. We compute them in closed form from per-branch taxon-weight summaries. For each branch *l* ∈ {*i, j, k*} of *u*, we define the following notations.

- 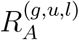 and 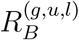: the sets of real taxa in the current subproblem that belong to partitions *A* and *B*, respectively, and are present in branch *l*.
- 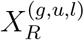 : the subset of real taxa represented by dummy taxon *X* that are present in branch *l*.
- 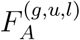 and 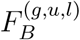: the full sets of real taxa in partitions *A* and *B*, respectively, within branch *l*. For example,

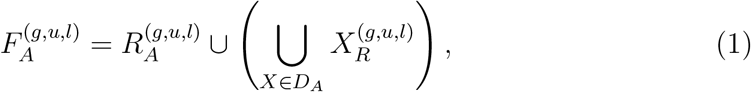

where *D*_*A*_ is the set of dummy taxa in partition *A*. The set 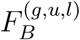 is defined analogously using 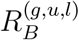 and *D*_*B*_.

For the child branches *i* and *j*, these sets are obtained from the descendant taxa of the corresponding branches. But quite importantly, For the parent branch *k*, only taxa present in the sibling branches of speciation nodes on the path from *u* to the root are included, since only those taxa can form SQs having anchor LCA *u*.

We now compute weights for the terms where the two taxa come from different branches of *u*. For two distinct regions *l, m* ∈ {*i, j, k*}, the set 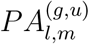 contains all unique unordered pairs *a, b* such that *a, b* ∈ *F*_*A*_, one taxon is present in region *l*, the other is present in region *m*, and the two taxa are not represented by the same dummy taxon in the current subproblem. Since the two regions are distinct, the raw product 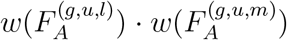 counts all candidate pairs between the two regions. However, it also includes invalid pairs in which both taxa are represented by the same dummy taxon. Therefore, using the inclusion-exclusion principle, we compute

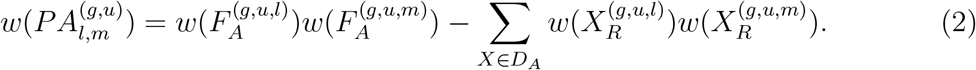

The corresponding quantity 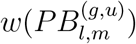 is computed analogously by replacing *F*_*A*_ and *D*_*A*_ with *F*_*B*_ and *D*_*B*_. Since the pairs are unordered, 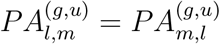 and 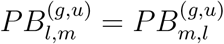.

Next, we compute taxon pair weights where both of the taxa come from the same branch. For a child branch *l* ∈ {*i, j*}, the set 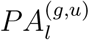 contains all unique unordered pairs *a, b* such that 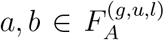 and the two taxa are not represented by the same dummy taxon in the current subproblem. The raw product 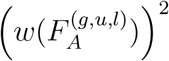 counts ordered pairs, including self-pairs and invalid same-dummy-taxon pairs. To obtain the valid unordered pairs, we subtract these invalid contributions and divide by two:

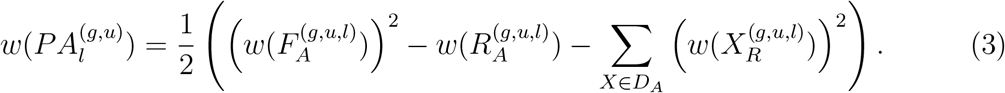

The quantity 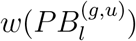 is computed analogously. These taxon pair weights can be computed for all speciation nodes using post-order tree traversals and cached taxon-weight information. For each speciation node, the summaries for its child branches are obtained from the taxa descending from those branches. The summary for the eligible parent-side region is obtained by combining the relevant sibling branches along the path from that node to the root. Thus, the pair weights required by the above formulas can be computed directly from the gene family tree, without explicitly enumerating quartet representatives.

##### Gain Calculation and Incremental Bookkeeping

Following the *Fiduccia-Mattheyses* (FM) heuristic used in wQFM-GDL, we iteratively improve the initial bipartition by transferring taxa from one partition to the other in order to find the bipartition which maximizes the quartet score. The heuristic may require multiple iterations, each consisting of several taxa transfers. Further details are provided in (38). Within the QFM framework, the *gain* of a taxon transfer is defined as the resulting change in the bipartition score due to that transfer. For a transfer of taxon *a* ∈ *A*, the gain is defined as:

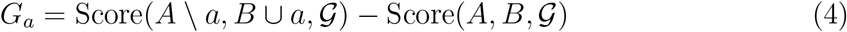

In a single iteration, multiple transfers take place and based on the cumulative gain, last few transfers might be rolled back. Now, importantly, naively calculating the bipartition scores separately and then using them to calculate gain would be quite inefficient. Thus, efficient gain calculation is quite important for the scalability of wQFM-GDL which requires some important observations. Thus, we employ some effective strategies to speed up the gain calculation, specially for the transfers of real taxa. Details are presented in Supplementary Section 2.1.

#### 2.3.4 Locus-aware normalization scheme

The prior normalization scheme used in wQFM-TREE could also be applied directly to wQFM-GDL. However, we identified an opportunity to improve normalization by introducing different normalization factors for distinct locus-specific regions in multicopy gene trees. Here, the concept of locus tree plays a crucial role. Rasmussen and Kellis proposed the DLCoal model (37) for jointly modelling GDL and ILS, which introduces a third tree called the locus tree, alongside the gene tree and the species tree. The locus tree is derived from the species tree via a top-down duplication–loss process, with each internal node labeled as either a speciation or a duplication event. A duplication event creates a new daughter locus from the parent locus. Both the parent and daughter loci evolve independently thereafter and can undergo further duplications or losses in subsequent evolutionary steps. Finally, a gene family tree is constructed from the locus tree by applying a multi-locus coalescent process.

Unlike the case with single-copy gene trees, a particular set of four species (e.g., *a, b, c, d*) can appear multiple times within different locus-specific subtrees of a particular gene tree. Now, note that the set of taxa in the various locus-specific subtrees may not be identical because each locus undergoes a distinct evolutionary history, marked by different duplication and loss events. Importantly, the normalization factors used in our method depends on the taxa set. The normalization scheme used in QMC and QFM families of methods does not consider this and thus any quartet on *a, b, c, d* in a gene tree is assigned the same weight. However, our observation is that, the quartets on a particular set of four species across different locus-specific subtrees should be assigned different weights as the taxa set of different locus-specific subtrees may vary. Recently, it has been shown that normalization of TREE-QMC can be affected in case of missing taxa in incomplete gene trees and thus needs correction (16). However, in case of gene family trees, identifying the taxa set associated with a particular loci within a multicopy gene tree is not straightforward. Our key step is that all quartets within an equivalence class descend from the same ancestral locus at the time of the speciation event corresponding to the anchor LCA, and the normalization factors for quartets associated with each anchor LCA are computed separately. Thus, each quartet equivalence class on *a, b, c, d* will have separate normalization factors and for all quartet equivalence classes associated with a particular anchor LCA, we compute normalization factors separately and independently. The variation in the normalization factors across the gene tree is primarily because of the different effective taxa set due to different duplication and loss history across the loci in the gene tree. To determine the effective taxa set relevant for a group of quartets, we first identify the subtrees that can contribute to forming them. For a speciation node acting as the anchor LCA, the taxa set includes (i) the taxa in the subtrees rooted at that node and (ii) the taxa descendant to the branches of the speciation nodes along the path from the anchor LCA to the root. Taxa outside these regions cannot participate in forming the corresponding quartets and are excluded. The resulting restricted taxa set is then used to compute normalization factors specific to that group of quartets (Figure 4). This locus-aware normalization scheme provides a substantial accuracy improvement for wQFM-GDL while incurring only a minimal runtime overhead.

**Figure 4:**
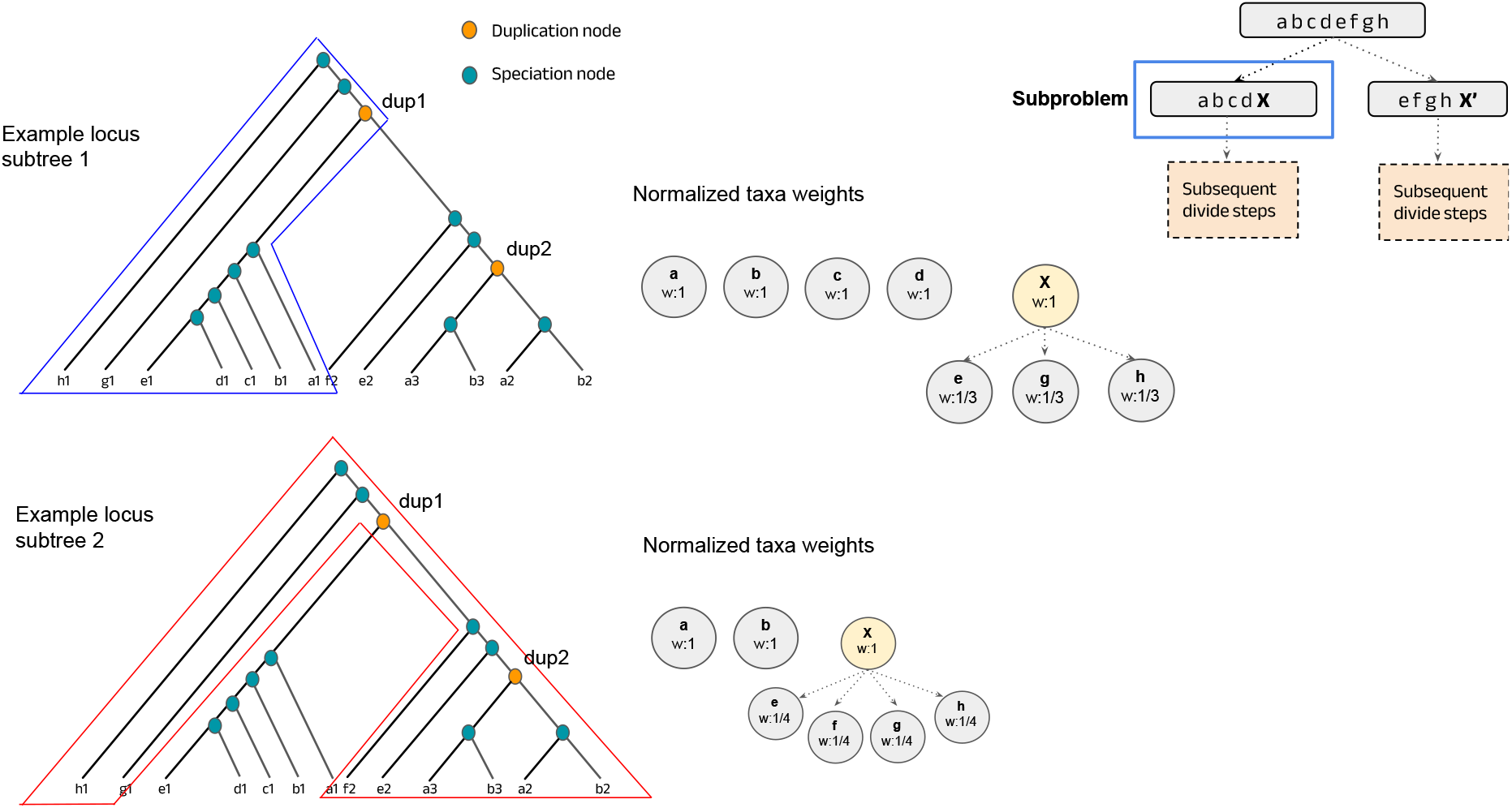
Different normalized taxon weights for quartets associated with different regions of the gene family tree. All quartets with anchor LCA in the blue region are restricted to taxa set *a, b, c, d, e, g, h* as *f* taxon is absent due to a loss event. Similarly, quartets with anchor LCA in the red region are restricted to the taxa set *a, b, e, f, g, h*. Thus, the normalization factors vary across the gene family tree.

It is important to note that this is an approximate strategy rather than an exact formulation, since the normalization factors of SQ equivalence classes sharing the same anchor LCA may still differ. However, our goal is to reduce the error in the traditional normalization scheme while keeping the computation tractable. Although more sophisticated approaches could be designed, our experimental observations suggest that they would likely increase time complexity without providing meaningful gains in accuracy.

We show an example in Figure 4. Here, duplication in the *dup*1 node creates a new locus, and both evolve in parallel and independently. In case of one of the loci that evolves along the left subtrees, *f* taxon is absent due to a loss event. All quartets with anchor LCA in the blue region are restricted to taxa set *a, b, c, d, e, g, h* as there is no *f* taxon, which results in the normalized taxon weights as shown in the Figure. On the other hand, in the right subtree, *c, d* is absent. Moreover, there is another duplication event at the node *dup*2. For quartets with anchor LCA in that right subtree, the effective taxa set is *a, b, e, f, g, h*. Thus, the normalization factors should be changed. Thus, unlike the traditional normalization strategy illustrated in Supplementary Figure 2, we employ multiple normalization factors for the same subproblem. The performance gain due to the locus-aware normalization scheme may vary based on the amount of duplication and loss in the dataset. We have discussed this in Section 3.3.4.

#### 2.3.5 Statistical Consistency and Local Support

wQFM-GDL solves the same optimization problem as ASTRAL-Pro, the Maximum per-Locus Quartet-score Species Tree (MLQST) problem. The exact solution to MLQST problem has been proven statistically consistent under the birth–death model called GDL developed by Arvestad et. al. (1), which only consider GDL in the absence of ILS, and under the MSC model which considers only ILS, in the absence of gene duplication and loss. However, there is no consistency proof for a model which combines ILS and GDL, such as DLCoal (37). wQFM-GDL serves as a heuristic for the MLQST problem.

We have added local posterior probability and branch length calculation for the species trees inferred by wQFM-GDL using the same approach used in ASTRAL-Pro, which is an intuitive and logical method in multicopy setting. For each internal branch of the estimated species tree, the branch defines a quadripartition of the taxa, corresponding to the topology displayed by that branch and its two alternative resolutions. The per-locus speciation-quartet counts supporting each of these three resolutions are computed and then used as input to the local posterior probability calculation process defined in (41).

### 2.4 wQFM-GDL-Q: Extending wQFM for GDL

wQFM takes as input a set of weighted quartets where the weight is the frequency of the quartet in the input gene trees. We devise an algorithm to create an appropriate weighted quartet set that only contains SQs and only one representative from an equivalence class. Then, this quartet set is given input to the wQFM algorithm which remains unchanged. We call this method wQFM-GDL-Q. For generating this quartet set, we adopt a similar algorithmic approach to the score calculation process in wQFM-GDL-T; however, in this case, we explicitly enumerate quartets with appropriate weights, rather than computing the sum of quartet weights. For each speciation node in the tagged and rooted gene trees, we independently generate the speciation-driven quartets with anchor LCA at that particular node and add them to the set. Similar to wQFM, the weight of a quartet is defined as its frequency across the gene trees. However, because the gene trees here are multicopy, multiple instances of the same quartet may occur within a single gene tree, and consequently, the total frequency (weight) of a quartet can be substantially larger than the number of gene trees, unlike in wQFM. An important point to note is that, although the high-level idea resembles the score calculation procedure of wQFM-GDL-T, there is no notion of satisfied or violated quartets here because there is no bipartition to consider at this stage. Instead, our objective is to generate all speciation-driven quartets (SQs) from the gene trees so that wQFM can operate on them. Since this step focuses on explicit quartet enumeration rather than efficiently computing scores directly from the gene trees, the procedure is more straightforward compared to wQFM-GDL-T. The algorithm for generating the quartet set is shown in Supplementary Algorithm 1.

As expected, wQFM-GDL-Q is considerably slower than wQFM-GDL-T. However, its main advantage is that under smaller model conditions, where explicit quartet enumeration is feasible, we observe that it outperforms competing methods, including wQFM-GDL-T, in several settings. This may be because the heuristics used in wQFM-GDL-T, which allow it to operate directly on gene trees, can sometimes influence its accuracy. Notably, the generated quartet set can be given input to any quartet amalgamation method. We compare the performance of the QMC framework with wQFM-GDl-Q in Section 3.3.1.

### 2.5 Complexity Analysis

The total time complexity for wQFM-GDL-T is *O*(*n*^2^*k* + *αn*^2^*Dd*), where *n* is the number of taxa, *k* is the number of gene trees, *D* is the number of unique speciation-driven tripartitions, *d* is the number of dummy taxa, and *α* is the number of refinement iterations (consistently observed to be ≤ 5 in our studies). It has successfully analyzed 500 taxa datasets with high duplication rate (more than 2,000 leaves) within 20 hours and 16GB of memory. The running time and memory consumption statistics for the methods used in this study are reported in Supplementary Section 4.

#### 2.5.1 Complexity of wQFM-GDL-T

We establish the time and memory complexity of wQFM-GDL below.

##### Time Complexity

###### Theorem 2.

*The time complexity of processing a single subproblem in wQFM-GDL-T is O*(*αnDd*), *where n is the number of taxa, D is the number of unique speciation-driven tripartitions, d is the number of dummy taxa in the subproblem, and α is the number of FM iterations*.

*Proof*. The time complexity of wQFM-GDL-T is determined by an iterative bipartition refinement process within a divide-and-conquer framework. To optimize the calculation of quartet scores, the algorithm identifies and stores the set of unique tripartitions associated with internal nodes tagged as speciation events across the input gene family trees. Let *D* denote the number of these unique speciation-driven tripartitions. By performing computations on this condensed set, the complexity of scoring and gain refinement is governed by *D* rather than the total size of the input gene trees.

The algorithm begins with a one-time construction of a greedy consensus tree from the input gene trees, which requires *O*(*n*^2^*k*) time for *k* gene trees on *n* taxa. In each sub-problem of the divide-and-conquer process, an initial bipartition is generated by scoring candidate bipartitions induced by the *O*(*n*) edges of this consensus tree. For each edge, the algorithm computes the score of the candidate bipartition in *O*(*dD*) time, where *d* is the number of dummy taxa in the subproblem. Consequently, selecting the best initial bipartition for a subproblem requires *O*(*nDd*) time.

Following the initial selection, the bipartition is refined using the Fiduccia-Mattheyses (FM) algorithm. Each iteration of the FM algorithm involves transferring (*r* + *d*) taxa between partitions, where *r* is the number of real taxa. Every transfer necessitates a gain recalculation that is dominated by the complexity of updating dummy taxon weights, taking *O*(*Dd*) time. Assuming the number of FM iterations is bounded by an empirical parameter *α*, the refinement process for a subproblem takes *O*(*α*(*r* + *d*)*Dd*) time. By adding the cost of finding the best initial bipartition to the cost of its iterative improvement, the total complexity of handling a single subproblem is *O*(*nDd* + *α*(*r* + *d*)*Dd*). Given that (*r* + *d*) is *O*(*n*), this total complexity is effectively *O*(*nDd* + *αnDd*), which simplifies to *O*(*αnDd*).

###### Corollary 1.

*Under balanced partitioning, the overall time complexity of wQFM-GDL-T is O*(*n*^2^*k* + *n*^2^*D* log *n*), *assuming α is a small constant*.

*Proof*. Since approximately *n* subproblems are required to construct the full species tree, the overall time complexity is *O*(*n*^2^*k* + *αn*^2^*Dd*). Under the assumption of balanced partitioning, where each divide step splits the taxa evenly, the number of dummy taxa in any subproblem is limited to *d* = *O*(log *n*). This leads to a total complexity of *O*(*n*^2^*k* + *αn*^2^*D* log *n*), which simplifies to *O*(*n*^2^*k* + *n*^2^*D* log *n*) for most practical applications as *α* is empirically found to be a very small constant (*α* ≤ 5).

###### Corollary 2.

*Under unbalanced partitioning (rooted caterpillar), the overall time complexity of wQFM-GDL-T is O*(*n*^2^*k* + *n*^3^*D*), *assuming α is a small constant*.

*Proof*. In cases where the bipartitions are unbalanced, *d* approaches *O*(*n*). Thus, the total complexity becomes *O*(*n*^2^*k* + *αn*^3^*D*), effectively *O*(*n*^2^*k* + *n*^3^*D*).

##### Memory Complexity

###### Theorem 3.

*The memory complexity of processing a single subproblem in wQFM-GDL-T is O*(*Dd*), *where D is the number of unique speciation-driven tripartitions in the gene family trees and d is the number of dummy taxa in the subproblem. Therefore, the memory complexity is O*(*D* log *n*) *under balanced partitioning and O*(*Dn*) *in the worst case*.

*Proof*. The memory complexity of wQFM-GDL-T is primarily driven by the storage of speciation-driven tripartitions and the auxiliary data structures required for efficient gain updates during the FM heuristic. To minimize redundant computations, the algorithm maintains the set of *D* unique tripartitions derived from the input gene family trees. For a given subproblem, the algorithm must track the relationship between these tripartitions and the *d* dummy taxa that represent the sister partitions in the divide-and-conquer tree. Specifically, for each unique tripartition, the algorithm stores weights and counts associated with the current distribution of dummy taxa to enable *O*(*Dd*) gain updates.

This results in a memory requirement of *O*(*Dd*) per subproblem. The overall memory complexity is not obtained by summing this quantity over all sub-problems, because wQFM-GDL-T processes subproblems sequentially. After a subproblem is processed, its auxiliary gain-update structures can be discarded before processing the next subproblem. Thus, the peak memory usage is determined by the subproblem containing the largest number of dummy taxa. If *d*_max_ = max_S_ *d*(S), where the maximum is taken over all subproblems S generated during the divide-and-conquer recursion, then the overall memory complexity is *O*(*Dd*_max_).

Since *d* is bounded by *O*(log *n*) under balanced partitioning, the memory complexity becomes *O*(*D* log *n*) in this case. In the worst case, where *d* can approach *O*(*n*), the memory complexity becomes *O*(*Dn*). Thus, the space complexity remains highly scalable for large phylogenomic datasets. This efficient use of memory allows wQFM-GDL-T to process thousands of gene family trees without the prohibitive overhead of storing full quartet sets.

#### 2.5.2 Complexity of wQFM-GDL-Q

##### Time Complexity

wQFM-GDL-Q follows a two-stage process: the explicit generation of a speciation-driven quartet set and the subsequent amalgamation of those quartets into a species tree using the original wQFM.

In the first stage, the algorithm iterates through *k* gene family trees to identify speciation-driven quartet (SQ) equivalence classes. Let *L* denote the maximum number of leaves, or gene copies, in any input gene family tree, and let *n* denote the number of distinct species. For single-copy gene trees, *L* ≤ *n*, whereas under gene duplication, *L* can exceed *n* depending on the number of duplicated copies per species. For each speciation node *s*, it performs list operations to gather taxa from subtrees and parent lineages. Listing real taxa in subtrees and generating all possible pairs takes *O*(*n*^2^) per speciation node in the worst case. Since a gene family tree with *L* leaves has *O*(*L*) internal nodes, this gives a worst-case cost of *O*(*Ln*^2^) per gene family tree. Across all *k* gene trees, the complexity of generating the refined set of *m* unique SQ equivalence classes is *O*(*kLn*^2^ + *m*). The *m* term accounts for storing or outputting the generated SQ equivalence-class representatives.

In the second stage, these *m* quartets are provided as input to wQFM. Based on the QFM Fast and Improved (QFM-FI) (**?**) complexity analysis, the amalgamation phase takes *O*(max(*nm*^2^, *n*^3^*m*)) time, where *n* is the number of distinct species. Consequently, the total time complexity for wQFM-GDL-Q is:

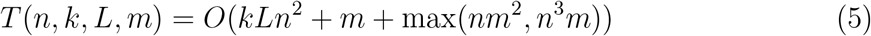

Since the amalgamation term already depends on *m*, this can also be written as:

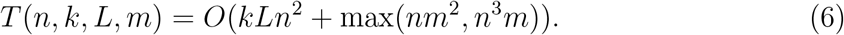

Given that the number of equivalence classes *m* can be as large as *O*(*n*^4^), and the value of *L* may also increase with the amount of duplication in the gene family trees, wQFM-GDL-Q is computationally more intensive than wQFM-GDL-T and is only suited for smaller datasets where explicit quartet generation is feasible.

##### Memory Complexity

The space complexity for wQFM-GDL-Q is determined by the storage of the explicit quartet set generated in the first stage. This requires *O*(*m* + *n*) memory to hold the *m* speciation-driven quartets and the taxon names.

## 3 Experimental results and discussions

### 3.1 Methods and measurements

We test the performance of wQFM-GDL-T and wQFM-GDL-Q on existing and our new simulated datasets and biological datasets against the leading methods: ASTRAL-Pro3 (50), SpeciesRax (30), FastMulRFS (29), and DupLoss-2 (33). On the simulated datasets, we compared the estimated trees with the model species tree using normalized Robinson–Foulds (RF) distance (39). We analyzed multiple data replicates under various model conditions and conducted a two-sided Wilcoxon signed-rank test (48) to compare the distribution of RF rates across different replicates in a particular model condition. We set the threshold *α* = 0.05 to measure the statistical significance of the differences between two methods. For the biological dataset, we compared the estimated species trees to established evolutionary relationships. We assessed support values in the estimated trees using local posterior probabilities (41) and quartet support (defined to be the number of quartets from the set of input gene trees that agree with the candidate species tree).

### 3.2 Datasets

#### 3.2.1 Simulated Datasets

We use S25 from the ASTRAL-Pro study (51), which contains 25 species under diverse duplication, loss, and ILS conditions, and S100, simulated by Molloy and Warnow (29) based on a real fungal dataset. We note that the number of model conditions for large datasets was limited in ASTRAL-Pro study. Therefore, to extensively evaluate the performance under GDL on large datasets, we generated two new large-scale simulated datasets comprising 200 and 500 taxa, which we call SIM200 and SIM500, respectively. In total, we analyzed 156 model conditions, including 72 large model conditions with 200 and 500 taxa, which was absent in the ASTRAL-Pro study. These include high-duplication settings with gene family trees containing up to 2,000 leaves.

For both SIM200 and SIM500, species trees and gene family trees were simulated using SimPhy (27) under varying duplication rates, loss rates, and ILS levels. Sequence alignments were generated using AliSim (23), and maximum likelihood gene trees were estimated with FastTree (35) under the GTR+Gamma model. Duplication rates and haploid effective population sizes were determined empirically. Detailed parameter settings and the full simulation pipeline are described in Supplementary Section 3.

#### 3.2.2 Biological datasets

We analyzed three important empirical datasets with multicopy gene trees (summarized in Table 1), Plants83 (46), Vertebrates188 (30), and Archaea364 (11), which cover a wide range of organisms. The vertebrates dataset is the largest dataset we analyzed, comprising 188 species and more than 30,000 gene families. The Archaea364 dataset is particularly challenging due to the presence of host–symbiont gene transfers in widely conserved marker genes.

**Table 1:**
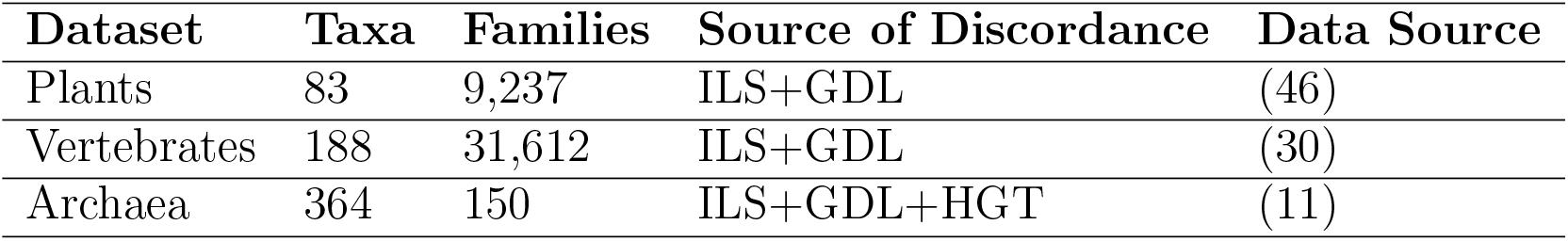
Empirical datasets used in our study.

| Dataset | Taxa | Families | Source of Discordance | Data Source |
| --- | --- | --- | --- | --- |
| Plants | 83 | 9,237 | ILS+GDL | (46) |
| Vertebrates | 188 | 31,612 | ILS+GDL | (30) |
| Archaea | 364 | 150 | ILS+GDL+HGT | (11) |

### 3.3 Results on simulated datasets

#### 3.3.1 Results for S25 dataset

In S25, both versions of wQFM-GDL and ASTRAL-Pro3 substantially outperform Species-Rax, DupLoss-2 and FastMulRFS across nearly all model conditions (Figure 5). We exclude DupLoss-2 and FastMulRFS for improved visual representation of the results because of their very high error rate, but results including DupLoss-2 and FastMulRFS are provided in Supplementary Section 5.2. While wQFM-GDL-T and ASTRAL-Pro3 perform similarly, wQFM-GDL-Q demonstrates better performance. It outperforms ASTRAL-Pro3 in 35 out of 52 model conditions, with statistically significant differences in 7 of these cases, and ties occurring in 8 conditions. As expected, all of the methods perform better with 500bp gene trees (lower gene tree estimation error) and lower ILS. Notably, as the duplication rate increases, all methods improve in performance, which is consistent with the findings of (51). This is likely because additional gene copies provide more information, similar to the benefit obtained from increasing the number of genes. It is important to note that SpeciesRax and DupLoss-2 are primarily designed to address GDL only, whereas the other evaluated methods can accommodate both ILS and GDL. Therefore, the datasets used in this study, which include both ILS and GDL, are not fully ideal for these methods. Nevertheless, both methods perform reasonably well on datasets with low levels of ILS. However, datasets that include both ILS and GDL are more realistic and biologically relevant. Therefore, the strong performance of wQFM-GDL and ASTRAL-Pro on these datasets further highlights their robustness and practical utility. Nevertheless, we included SpeciesRax and DupLoss-2 methods to better understand their behaviour under such evolutionary scenarios.

**Figure 5:**
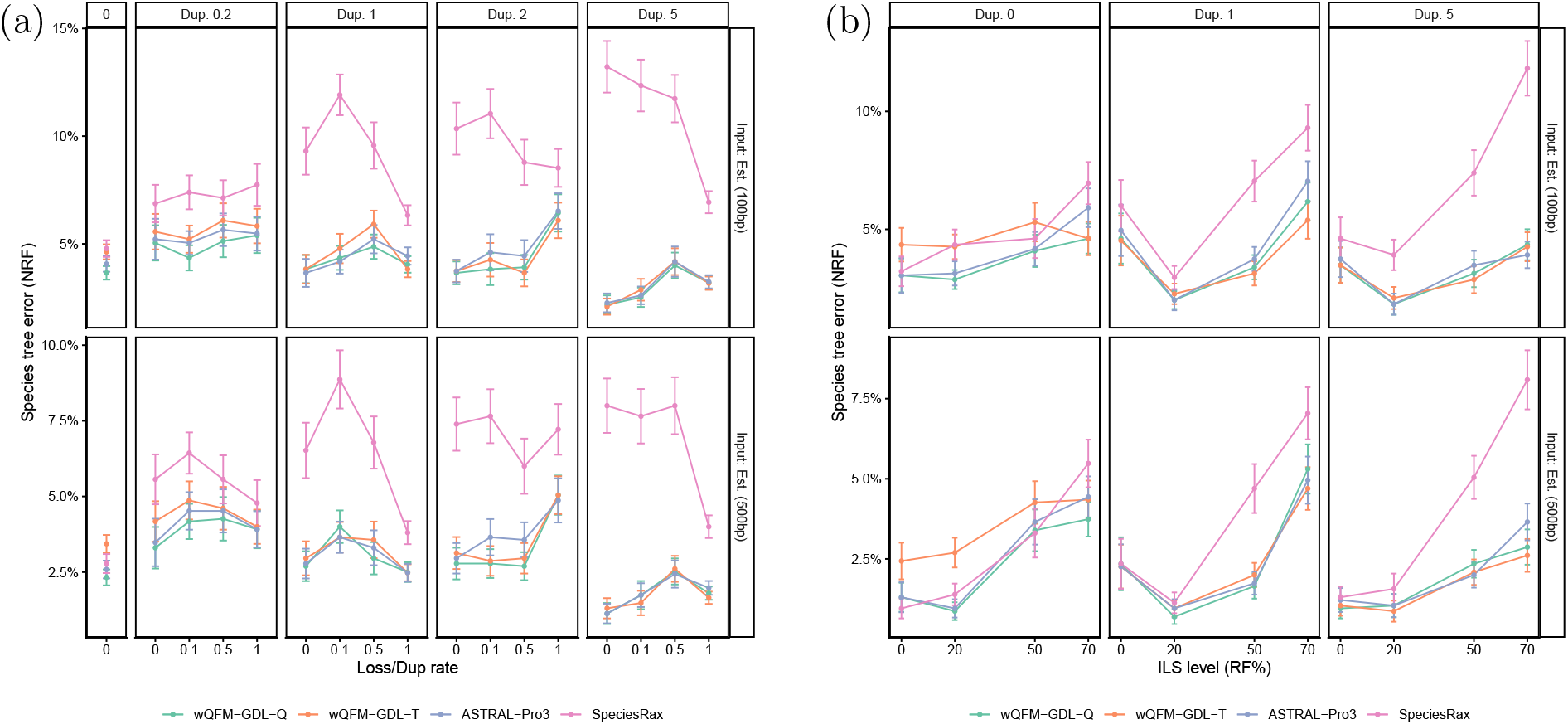
Species tree error on the S25 dataset using 1,000 estimated gene trees from 100 and 500bp alignments. We show the average RF rates with standard error bars over 50 replicates. (a) Controlling duplication rate (box columns, labelled by mean number of copies per species minus one) and loss rate (labelled by ratio of loss and duplication rate). (b) Controlling duplication rate and ILS level (RF rate between true gene trees and species tree). Results including DupLoss-2 and FastMulRFS are presented in Supplementary Section 5.2.

##### Comparison of Quartet Amalgamation Frameworks

In wQFM-GDL-Q, we first construct a quartet set containing only SQs, which is then given input to the wQFM algorithm. Now, the generated quartet set can be given input to any quartet amalgamation method, enabling it to process multi-copy gene trees and account for GDL. Thus, we test the performance of wQMC, a popular quartet amalgamation method, and compare it with wQFM-GDL-Q. This experiment also helps to understand the merit of QMC framework in this problem. The quartet file prepared with our algorithm is given input to both wQFM and wQMC. Figure 6 presents the performance of both algorithms on the S25 dataset. We observe that wQFM-GDL-Q is considerably better than wQMC on most of the model conditions. However, there is a recently developed scalable version of wQMC, TREE-QMC, which works directly on the gene trees and introduces new normalization techniques to improve results. TREE-QMC can work on multi-individual trees, but extending it with a duplication-aware quartet score for multicopy gene trees can be an interesting direction.

**Figure 6:**
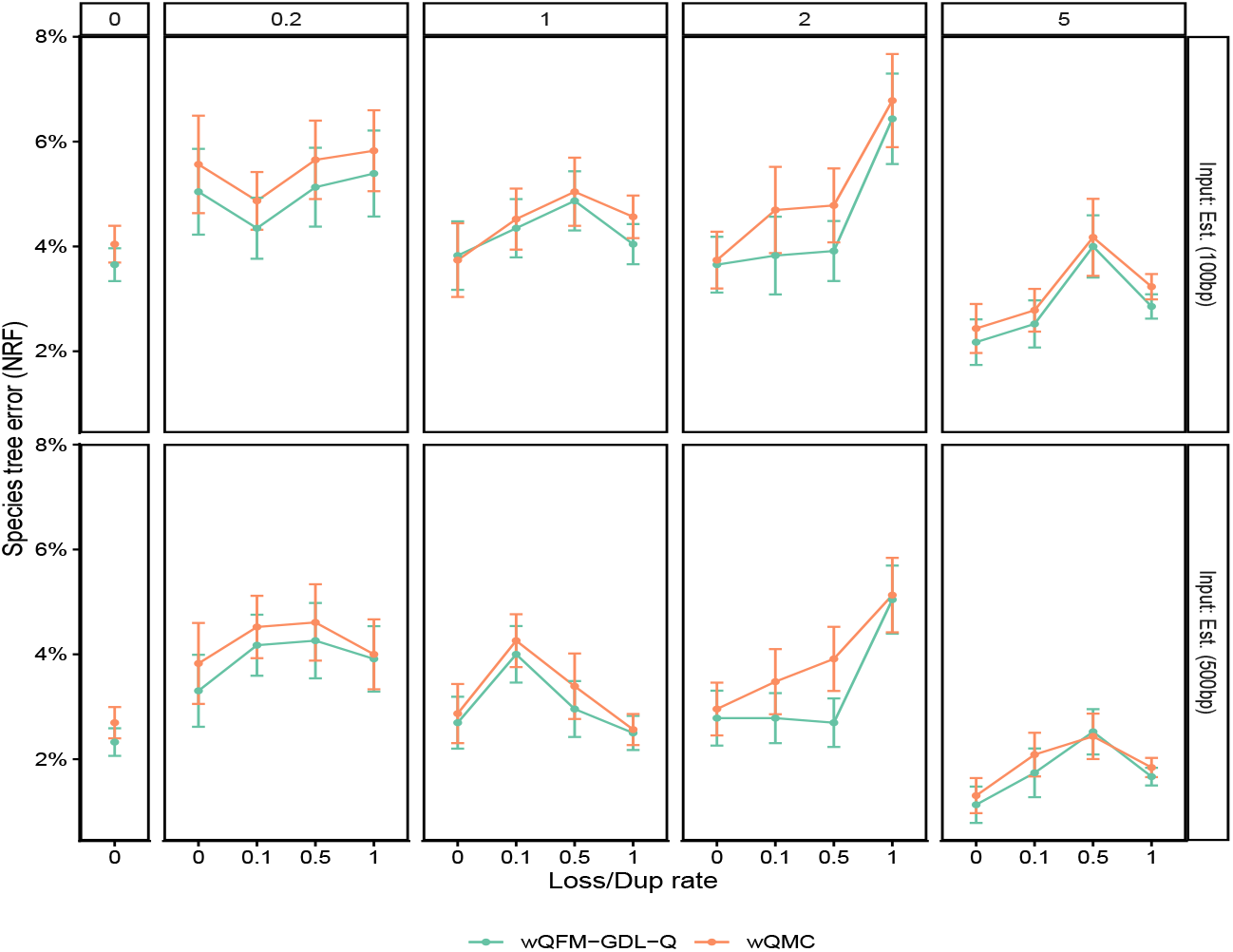
Species tree error on S25 dataset for wQFM-GDL-Q and wQMC (with a quartet file of only SQs using our algorithm).

#### 3.3.2 Results for S100 dataset

The performance of the methods on the S100 dataset is shown in Figure 7. Here, we can observe the accuracy of the methods with varying sequence length and Psize (effective haploid population size). Again, both versions of wQFM-GDL outperform ASTRAL-Pro and FastMulRFS in most of the model conditions, with wQFM-GDL-Q being the slightly more accurate version, although at 100 species, the quartet generation phase may take hours. We observe that, as the model conditions become more challenging with small sequences, such as in 25bp model conditions, wQFM-GDL performs noticeably better than the other methods. On the other hand, for the easier model conditions with larger 100bp sequences, the performances are quite similar. For both of the values of effective haploid population size, we observe similar trends.

**Figure 7:**
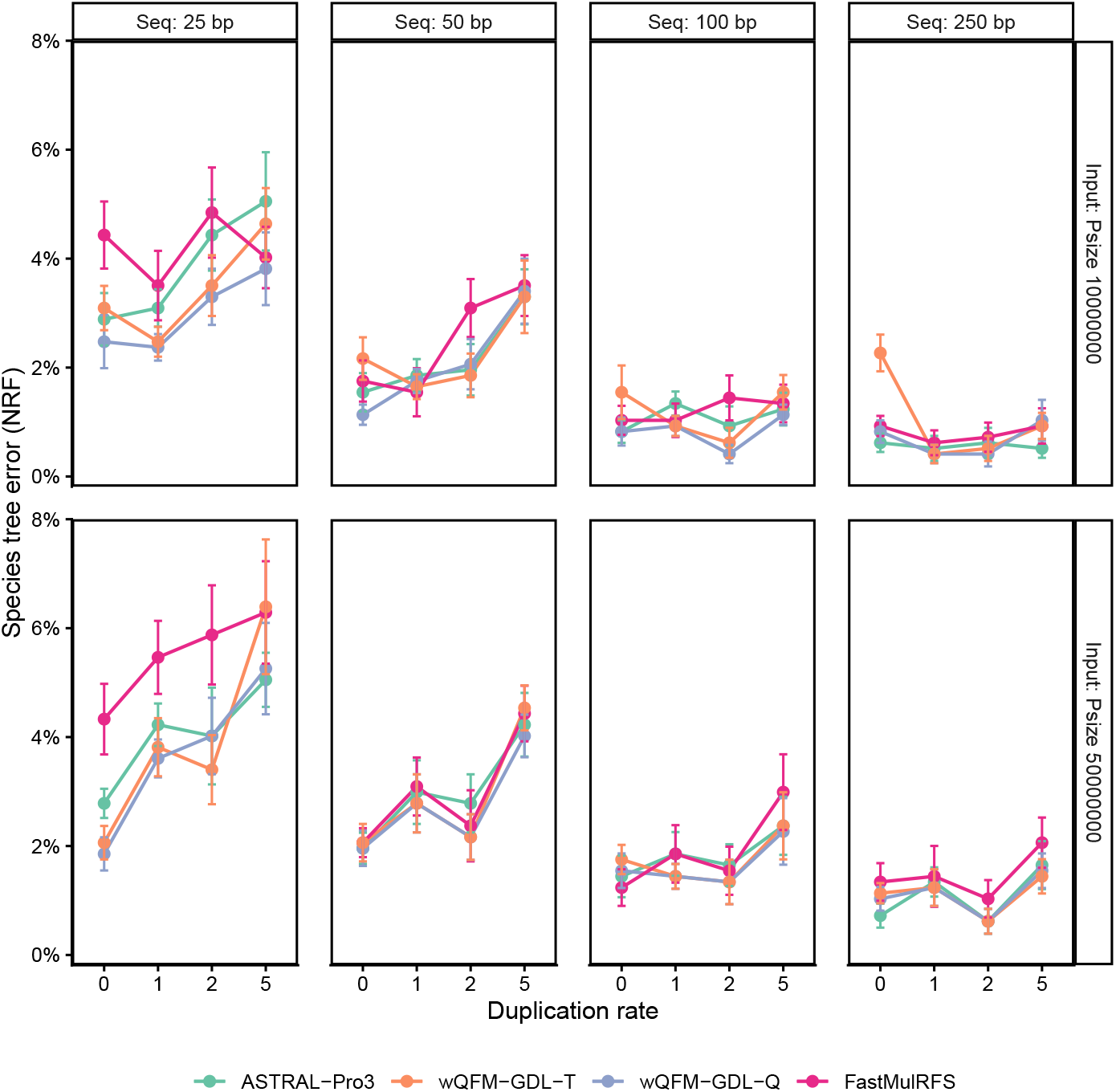
Species tree error on the S100 dataset for 100 ingroup taxa, 1000 gene trees, varying sequence length, and the effective haploid population size (labeled as Psize).

#### 3.3.3 Results for SIM200 and SIM500 datasets

Figure 8 presents the performance of the methods on SIM200 and SIM500 datasets. wQFM-GDL-Q could not be executed on these datasets due to its computationally intensive quartet generation. We could not finish running SpeciesRax on a single replicate with 500 or 1000 genes for some model conditions even after running for more than two days. Therefore, within our resource limits (64 GB of CPU RAM and 48 hours of computation per replicate), we could only analyze SpeciesRax on 250-gene model conditions of SIM200. Again, results including DupLoss-2 are in the Supplementary because of high error rate. Remarkably, wQFM-GDL-T convincingly outperformed all the methods in all 72 out of 72 model conditions, differences being statistically significant (*p <* 0.05). wQFM-GDL-T delivers almost 19% and 23% reduction in error compared to ASTRAL-Pro3 in SIM200 and SIM500, respectively. As expected, the performance of all methods decreases with fewer gene trees and higher levels of ILS.

**Figure 8:**
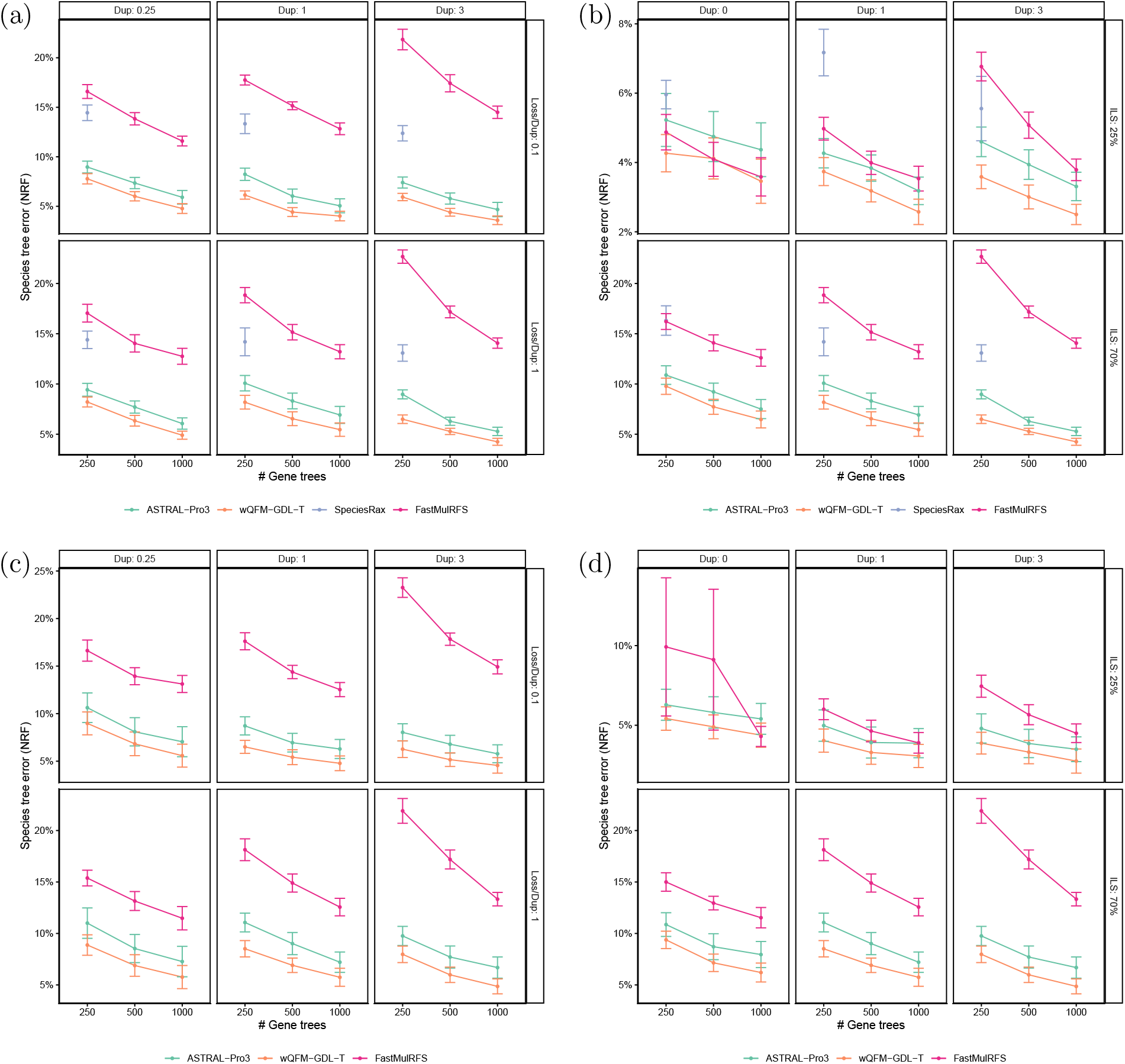
Species tree error on the SIM200 (a-b) and SIM500 (c-d) dataset using estimated gene trees from 100 bp alignments. (a,c) Controlling duplication and loss rate. (b,d) Controlling duplication rate and ILS level. SpeciesRax only completed 250-gene model conditions of SIM200 within our resource limits. Results including DupLoss-2 are presented in Supplementary Section 5.2.

The performance advantage of wQFM-GDL becomes more substantial as the model conditions increase in scale. The average error reductions over the closest competing method, ASTRAL-Pro3, increased from around 11% on S25 to 19% on SIM200 and 23% on SIM500. ASTRAL-Pro3 uses dynamic programming on a constrained search space, but wQFM-GDL uses the FM heuristic on a broader region of the full search space. The effectiveness of the inherent FM search strategy becomes more pronounced in larger search spaces, as observed in prior QFM works (36), and this trend appears to hold to the present setting with GDL as well. While wQFM-GDL leads to quite high accuracy, it is currently slower than some alternative methods. However, it remains scalable enough to analyze large datasets containing thousands of taxa and gene trees (Supplementary Section 4).

#### 3.3.4 Performance gain from the locus-aware normalization scheme

The performance improvement of the locus-aware normalization scheme on a few example model conditions of SIM200 and SIM500 is shown in Figure 9. As we can observe, when there is a considerable amount of duplication and loss, the differences among the normalization factors of the locus-specific subtrees are high and there is a noticeable performance boost in almost all model conditions, likely due to the significant differences in normalization factors among different loci (Figure 9a). However, when there is low duplication and loss, we observe little improvement (Figure 9b). Moreover, in model conditions with no duplication at all, the result remains the same.

**Figure 9:**
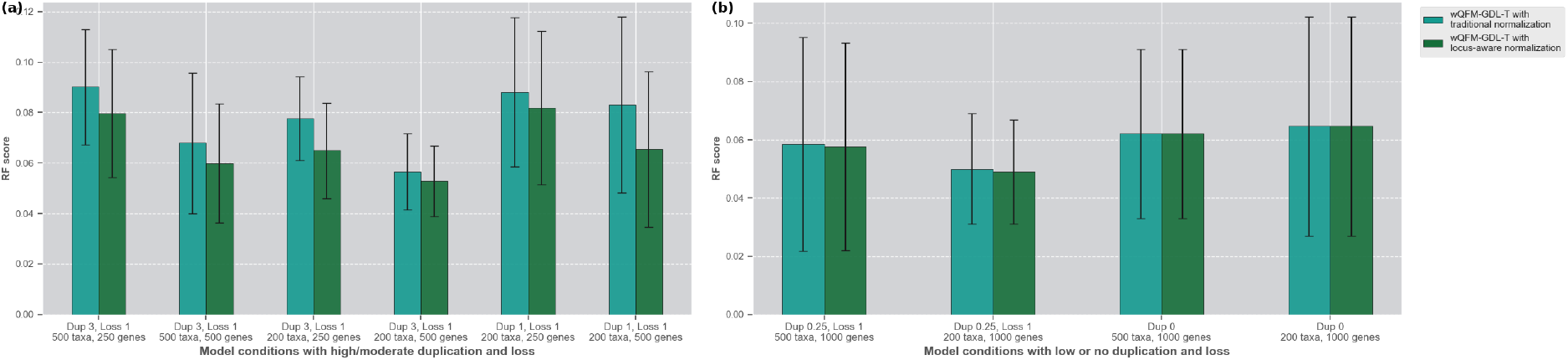
Comparison of the traditional and locus-aware normalization schemes across different model conditions of SIM200 and SIM500. (a) Significant performance gain in moderate to high duplication and loss scenarios. (b) Little to no performance gain in very low duplication and loss scenarios.

### 3.4 Results on biological datasets

#### Plants83 dataset

We analyze 9,237 multicopy gene trees of the Plants83 Dataset, which were inferred in the original study by Wicket et. al., (46). The tree estimated by wQFM-GDL (Figure 10) successfully recovers all major relationships and does not violate any known consensus among scientists. Following the most prominent recent methods, wQFM-GDL reconfirms the monophyly of Bryophytes (mosses, liverworts, and hornworts). Lycophytes are correctly recovered as the sister group of ferns and seed plants, and Gymnosperms are correctly recovered as the sister to flowering plants. It places Zygnematales (not Chara) as sister to all land plants and Amborella as sister to the rest of angiosperms. All of these relationships are well established. The trees inferred by wQFM-GDL, ASTRAL-Pro3, and SpeciesRax are largely identical, differing only in three contentious regions. In contrast, the tree inferred by DupLoss-2 differs substantially from those inferred by the other methods and conflicts with several well-established biological relationships. For example, DupLoss-2 recovers Lycophytes and Hornworts as sister groups, fails to place Amborella as sister to the remaining angiosperms, and shows additional discrepancies in several intra-clade relationships. FastMulRFS failed to complete the analysis on our machines, terminating with a memory allocation error across multiple attempts.

**Figure 10:**
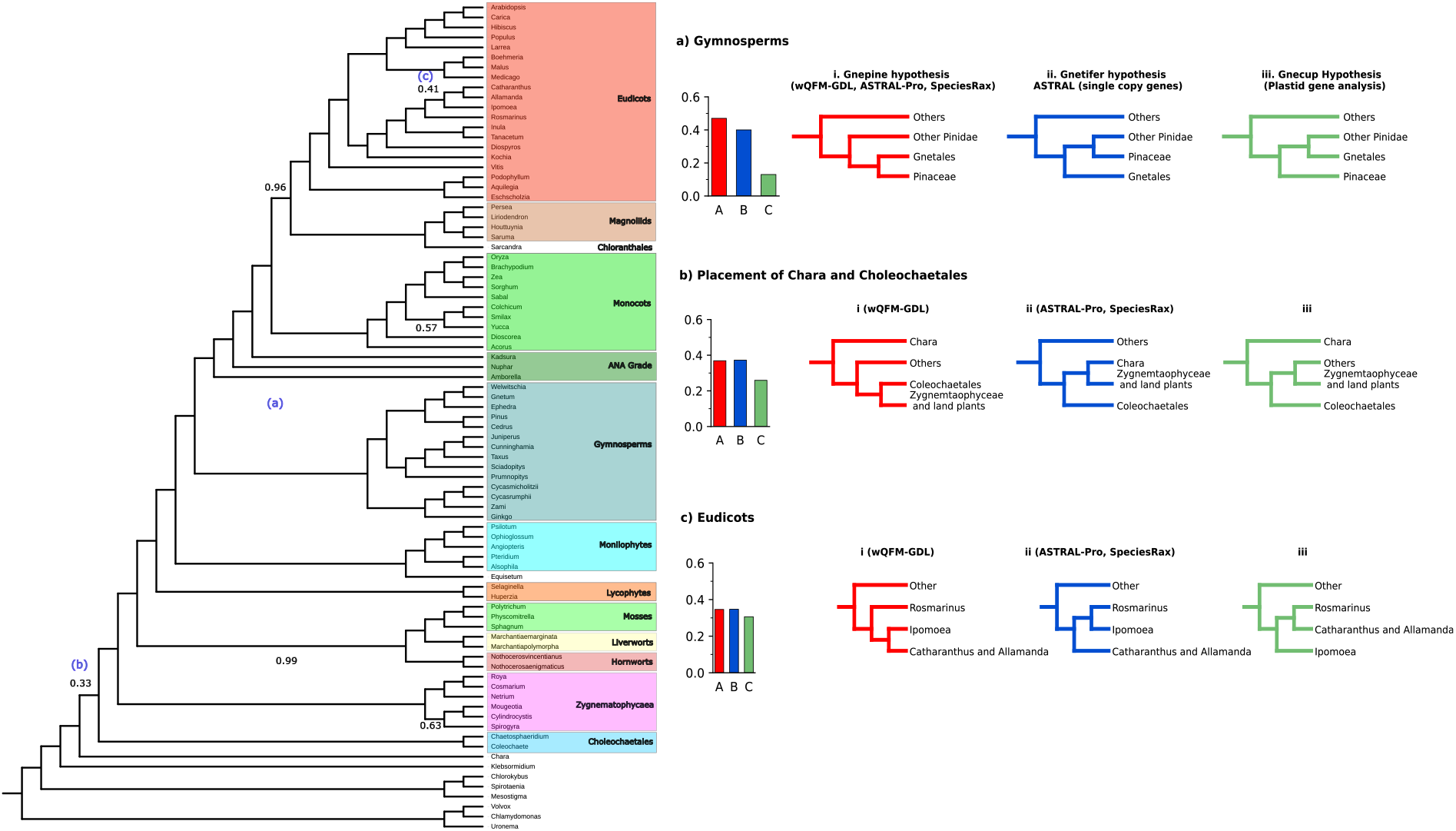
The species tree inferred using wQFM-GDL for the Plants83 dataset and alternative branching orders for contentious relationships.

The three quite contentious areas with alternative branching orders and quartet supports for those relationships are shown in Figure 10. Firstly, wQFM-GDL, ASTRAL-Pro3, and SpeciesRax all support the Gnepine hypothesis for Gymnosperms (1.0 local posterior probability), placing Gnetales and Pinaceae as sisters. However, in the 1kp plant analysis(47), ASTRAL with single-copy nuclear gene trees and plastome-based supermatrix analysis strongly suggested otherwise. Secondly, for the relative position of Coleochaetales and Chara, wQFM-GDL places Chara as the closest relative of Zygnematophyceae and Embryophyta (land plants), whereas ASTRAL-Pro3 and SpeciesRax place Coleochaetales algae. Both of the cases have almost exactly the same quartet support (36%). Finally, in the ASTRAL-Pro3 tree, Rosmarinus and Ipomoea are placed in the same clade. In contrast, wQFM-GDL recovers Ipomoea as sister to Catharanthus and Allamanda. For the second and third cases, the gene trees appear to contain nearly equal support for two alternatives, as shown in Figure 10. Therefore, additional evidence may be needed to resolve these relationships more confidently.

#### Vertebrates188 dataset

The tree inferred by wQFM-GDL-T is consistent with the NCBI taxonomy tree that is available on the Ensembl Compara database (12) except for only five branches. The other methods recover almost identical trees except these five branches, though DupLoss-2 differs from the others in a few more branches. FastMulRFS failed to complete the analysis on our machines because the available 64 GB of RAM was insufficient. All tested methods on this dataset violated a well-established phylogenetic relationship. The elephant shark, which is a *Chondrichtyian*, is placed within *Osteicthyians*, as sister to *Sarcopterygii*. wQFM-GDL also recovers the same relationship.

The other cases are quite contentious. In Platyrrhini (monkey sub order), when placing *Aotidae, Cebus/Saimiri*, and *Callitrichidae*, wQFM-GDL and ASTRAL-Pro group *Aotidate* with *Cebus/Saimiri* (Supplementary Figure 6). The taxonomy places *Cebus/Saimiri* and *Callitrichidae* together, while SpeciesRax and DupLoss-2 place *Aotidate* and *Callitrichidae* together. There exist studies that agree with SpeciesRax (34) and also wQFM-GDL and ASTRAL-Pro (40).

In case of the avian subtree of 24 species, the tree estimated by wQFM-GDL matches the recent bird phylogeny of 363 taxa published by Stiller et. al., (43). However, the taxonomy groups the *Estrildidae* and *Fringillidae* together, whereas all of the tested methods group *Fringillidae* and *Passerellidae* together. Other contentious cases include the placement of *Ambassidae*. All the tools place *Cichliformes* as sister to *Ambassidae* whereas the taxonomy places *Pomacentridae* as sister to *Ambassidae*. The full tree and some important contentious subtrees inferred by wQFM-GDL can be found in Supplementary Section 5.3.

#### Archaea364 dataset

DPANN archaea represent a potentially deep-branching, monophyletic lineage characterized by small cell sizes and genomes. The monophyly, early emergence, and evolutionary significance of the various DPANN clades remain subjects of debate. The main difficulty in resolving the archaeal tree is the presence of host–symbiont gene transfers in the broadly conserved marker genes, in which members of the host-associated DPANN Archaea are sometimes grouped with their hosts in the gene trees. The original study used concatenation with ML for their analysis and did not explore any summary method analysis. wQFM-GDL successfully reconstructed the species tree from the gene trees provided in (11) as shown in Figure 11. We used not only the full set (Supplementary Figure 7) but also the top 50% and 25% marker proteins (Figure 11), which filter out the incongruent or contaminated markers.

**Figure 11:**
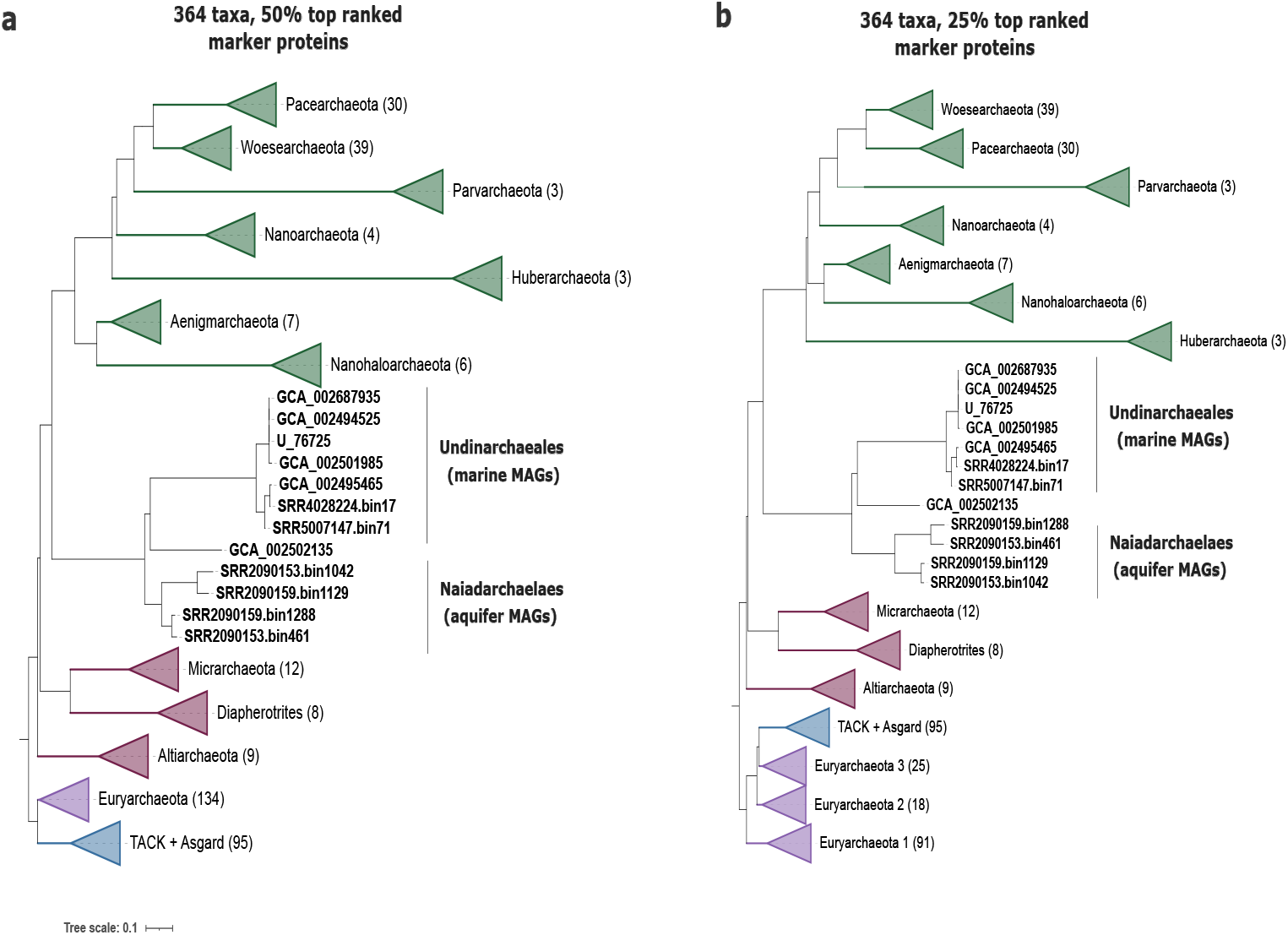
The species trees inferred with wQFM-GDL from the Archaea364 dataset with top 50% and 25% marker proteins.

Our analyses recovered DPANN archaea (including Undinarchaeota) as a monophyletic group as a whole. Furthermore, both the species trees from the top 50% and 25% marker proteins suggested that Undinarchaeota form a distinct lineage that branches between two other DPANN clans with maximum branch support. The first clan consists of Altiarchaeota, Micrarchaeota, and Diapherotrites, and the second clan consists of Woesearchaeota, Pacearchaeota, Parvarchaeota, Nanoarchaeota, Huberarchaeota, Aenigmarchaeota, and Nanohaloarchaeota. Undinarchaeota formed two orders, consisting of aquifer-derived and ocean-derived MAGs, i.e., the Naiadarchaeales and Undinarchaeales, respectively. All these findings match the findings of Dombrowski et. al., (11), done by CA-ML analysis, except for two slight dissimilarities associated with the position of Altiarchaeota inside the first clan and the relative position of Parvarchaeota and Nanoarchaeota in the second clan. Other methods also support the monophyly of DPANN and put Undinarchaeota between the two DPANN clans. However, there are some discrepancies in the tree inferred by DupLoss-2 and ASTRAL-Pro. ASTRAL-Pro puts Micrarchaeota and Diapherotrites in the second clan previously mentioned, with only Altiarchaeota in the first clan, which deviates from the findings of Dombrowski et. al. (Figures in Supplementary Section 5.3).

The tree reconstructed from the full set, top 50%, and top 25%, all showed similar findings. However, in the species tree of the top 25% marker proteins, the Euryarchaeota group is divided into multiple subgroups by all the algorithms.

## 4 Conclusions

In this study, we proposed wQFM-GDL, a highly accurate quartet-based summary method, which accounts for both ILS and GDL. We have leveraged the concept of SQs, utilized in the GDL model of ASTRAL-Pro within the QFM framework. Although we adapted the concept of speciation-driven quartets developed in ASTRAL-Pro, incorporating that into the divide-and-conquer framework and the Fiduccia–Mattheyses heuristic of the QFM family of methods required substantial algorithmic re-engineering. This includes the development of efficient techniques for constructing initial bipartitions in the QFM framework, as well as new combinatorial methods for computing refined quartet scores directly from gene family trees. We also introduce a locus-aware normalization scheme to improve accuracy.

wQFM-GDL is the first quartet amalgamation-based method that explicitly addresses GDL, and, after ASTRAL-Pro, only the second quartet-based method designed to handle both ILS and GDL. The main strength of wQFM-GDL, as observed from our experiments, is its high accuracy. In smaller or simpler model conditions, such as datasets with fewer taxa or longer sequences, wQFM-GDL performs on par with its main competitor, ASTRAL-Pro. However, in more difficult model conditions involving larger numbers of taxa, wQFM-GDL outperforms all competing methods, including ASTRAL-Pro, across all 72 out of 72 large-dataset model conditions. The results on empirical biological datasets are also promising, as wQFM-GDL recovers trees that do not violate any well-established biological relationships. Although wQFM-GDL is slower than methods such as ASTRAL-Pro and FastMulRFS, it remains reasonably scalable and can analyze datasets with hundreds of taxa and gene trees with high duplication and loss rates within hours.

In conditions involving both GDL and a significant amount of ILS, quartet-based methods such as ASTRAL-Pro3 and wQFM-GDL achieve the best performance and outperform the other methods considered in our study. Although ASTRAL-Pro and wQFM-GDL address the same optimization problem, the higher accuracy of wQFM-GDL compared to ASTRAL-Pro can likely be attributed to differences in the explored search space. ASTRAL-Pro uses exact dynamic programming within a constrained search space. In contrast, the FM heuristic used in wQFM-GDL methodically moves taxa between the two sides of candidate bipartitions, allowing the method to explore different regions of the solution space. This is also true in ILS-only condition, as we observed in our previous studies where wQFM/wQFM-TREE performed better than ASTRAL-III, specially in larger model conditions (26; 36). Moreover, the concept of satisfied and violated SQs used in wQFM-GDL, provides an effective measure of bipartition quality in this setting, which helps to find high-quality bipartitions of the species tree. These factors likely contribute to the strong empirical performance of the heuristic in practice.

Currently, wQFM-GDL uses the tagging and rooting heuristic introduced by ASTRAL-Pro. Complex evolutionary scenarios involving ILS and GDL may cause this heuristic to produce suboptimal rooting and tagging, as discussed in the paper, potentially reducing the accuracy of wQFM-GDL. Future works should explore designing a better tagging and rooting heuristic, which might enhance the performance of quartet-based methods. Furthermore, the divide-and-conquer structure of QFM naturally supports parallel execution, and a multi-processor-based implementation could significantly improve the scalability of QFM-based methods, including wQFM-GDL.

## Data availability

The simulated datasets generated for this study have been deposited in Dryad and are available at http://datadryad.org/share/LINK_NOT_FOR_PUBLICATION/DDMJOG0jvVsZvXHJQ1zmeMCHGrV0CNpV3PuikdMEeOQ. The remaining datasets analyzed in this study were obtained from previously published studies and are publicly available through the sources cited in the main text.

