## Supplementary information for "Highly Accurate Species Tree Inference from Orthologs and Paralogs through Scalable Quartet Amalgamation"

### List of Figures

1 Examples of speciation-driven quartets (SQs), Non-SQs, and quartet equivalence classes. . . . . . . . . . . . . . . . . . . . . . . . . . . . . 4

2 **Normalized taxon weights for different subproblems in the example divide phase of wQFM-TREE.** Real and dummy taxa in each subproblem are assigned a weight of 1, and the weights of the taxa represented by a dummy taxon are normalized so that their total sums to 1. For subproblem A, no dummy taxon is present, and normalization has no effect. For subproblem B, this is a simple case where the dummy taxon $X$ represents taxa 3 and 5. For subproblem C, we observe that the taxa represented by dummy taxon $Y$ are not assigned equal weights. Instead, they are weighted based on their relevance to the subproblem and subproblem structure. Since taxa 3 and 5 under dummy taxon $Y$ are not directly represented by $Y$, but rather through $X$, they receive lower weights compared to taxa 1 and 7. . . . . . . . . . . . . . . . . . . . . . . . . . . . . 9

3 **Different normalized taxon weights for quartets associated with different regions of the gene family tree.** All quartets with anchor LCA in the blue region are restricted to taxa set $a, b, c, d, e, g, h$ as $f$ taxon is absent due to a loss event. Similarly, quartets with anchor LCA in the red region are restricted to the taxa set $a, b, e, f, g, h$. Thus, the normalization factors vary within the gene family tree. . . . . . . . . . . . . . . . . . . . 11

4 Species tree error on the S25 dataset, including DupLoss-2 and FastMulRFS.(a) Controlling duplication rate (box columns, labelled by mean number of copies per species minus one) and loss rate (labelled by ratio of loss and duplication rate). (b) Controlling the duplication rate and ILS level (RF rate between true gene trees and species tree). . . . . . . . . . . . . 20

5 Species tree error on the SIM200 (a-b) and SIM500 (c-d), including DupLoss-2. (a,c) Controlling duplication and loss rate. (b,d) Controlling the duplication rate and ILS level. . . . . . . . . . . . . . . . . . . . . . . . . . 21

6 Important contentious subtrees of the species tree estimated by wQFM-GDL for the Vertebrates188 dataset . . . . . . . . . . . . . . . . . . . . . 22

7 The species tree inferred with wQFM-GDL for the Vertebrates188 dataset. 23

8 The species trees inferred with wQFM-GDL from the Archaea364 dataset with top 50% and 25% marker proteins . . . . . . . . . . . . . . . . . . . 24

9 The species tree inferred with wQFM-GDL for the Archaea364 dataset using the full set of protein markers. . . . . . . . . . . . . . . . . . . . . 25

10 The species tree inferred with ASTRAL-Pro3 for the Archaea364 dataset 26

### 1 Speciation-driven quartets and Equivalence class

Figure 1 illustrates examples of quartets in a gene tree that qualify as speciation quartets (SQs) and those that do not under the ASTRAL-Pro model. For a quartet to be considered SQ, the LCA of every combination of three leaves must correspond to a speciation node. Each quartet yields four such triplets, obtained by excluding one taxon at a time, and therefore four corresponding LCA nodes, though these need not be distinct. For the SQs shown in Figure 1, all LCAs are speciation nodes and thus carry a speciation signal, whereas for non-SQs, at least one LCA corresponds to a duplication node.

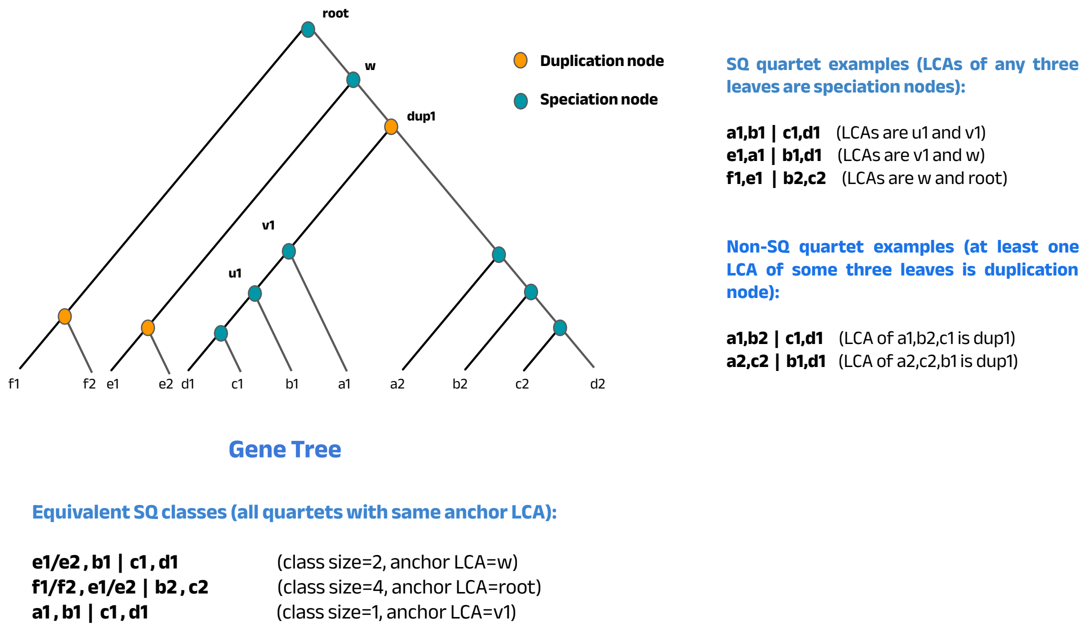

Figure 1: Examples of speciation-driven quartets (SQs), Non-SQs, and quartet equivalence classes.

To understand why this condition corresponds to having speciation information, let us consider a simple example. In the gene tree, a duplication event occurs at the node $dup1$ and two loci independently generate the quartets $a1, b1|c1, d1$ and $a2, b2|c2, d2$ through speciation events. These two quartets independently evolve through speciation, and both carry valid speciation information about the taxa $a, b, c$, and $d$. LCAs of every combination of three leaves for these quartets are speciation nodes. In contrast, consider the quartets $a1, b2|c1, d1$, $a2, b2|c1, d1$ etc. For these quartets, the LCA of some combinations of three leaves is $dup1$. We observe that the speciation information regarding $a, b, c, d$ in the two loci are already encoded in quartets $a1, b1|c1, d1$ and $a2, b2|c2, d2$ and the existence of these quartets are solely due to paralogy (duplication event at node $dup1$) rather than true speciation history.

Moreover, we have shown some examples of equivalent SQ classes. Anchor LCA is the LCA of the two anchor nodes of the quartet, and quartets on the same four species with the same anchor LCA are considered equivalent and one unit, as they carry the same speciation information.

For example, let us consider the quartets $e1, b1|c1, d1$ and $e2, b1|c1, d1$. Taxon $e$ is separated from $b, c, d$ in the speciation node $w$, and the relationship among $b, c, d$ in these two quartets is determined by the speciation event at $u1$. Thus, we are observing two quartets solely because of the duplication event at the MRCA of $e1$ and $e2$. These two quartets contain only one unit of speciation information about the taxa $b, c, d, e$. Thus, they should not be counted as two separate quartets and are part of the same equivalence class.

### 2 wQFM-GDL: Additional Details

#### 2.1 wQFM-GDL-T: Gain Calculation and Incremental Bookkeeping

The *Fiduccia-Mattheyses* (FM) heuristic in wQFM-GDL iteratively improves a bipartition by transferring taxa to the opposite partition to maximize the score. For a taxon $a$ currently in partition $A$, the gain $G_a$ is defined as the change in the total quartet score if $a$ were moved to partition $B$:

$$G_a = \text{Score}(A \setminus \{a\}, B \cup \{a\}, \mathcal{G}) - \text{Score}(A, B, \mathcal{G}) \tag{1}$$

Efficiently calculating these gains is critical to the algorithm's scalability.

##### 2.1.1 Efficient Gain Calculation for Real Taxa

We leverage the property that the change in the quartet score at a specific speciation node $u$ is identical for any real taxon $a \in R_A$ located within the same branch $l \in \{i, j, k\}$. Since every such real taxon contributes a unit weight and interacts with the rest of the bipartition symmetrically within the topology of $u$, the algorithm avoids redundant calculations:

1. We traverse the unique speciation tripartitions.
2. At each node $u$, we simulate a single real taxon transfer for each branch $l$ and compute the local change in satisfied and violated quartet weights ($\Delta sat$ and $\Delta vio$).

3. These local changes are stored for each branch and subsequently accumulated using a linear graph traversal to determine the total gain $G_a$ for all $a \in R_{A \cup B}$.

##### 2.1.2 Gain Calculation for Dummy Taxa

A real taxon in the current subproblem occupies a single, fixed position within a specific branch of a gene tree. In contrast, a dummy taxon $X_j$ serves as a representative for a set of real taxa from previous subproblems. The constituent real taxa of a single dummy taxon may be distributed across multiple branches of a speciation node $u$ depending on the gene tree's topology. As a result, transferring a dummy taxon does not produce a uniform score change across the set of dummy taxa at node $u$; instead, the impact on the score is unique to the specific distribution of that dummy taxon's underlying real taxa. Therefore, the gain for each dummy taxon must be calculated individually. For every unique speciation node $u$, the algorithm explicitly computes the local score change caused by the simulated transfer of $X_j$ and sums these differences across all $D$ tripartitions to determine the total gain.

##### 2.1.3 Incremental Bookkeeping of Pair Weights

A key optimization in wQFM-GDL is the localization of stateful bookkeeping variables at each internal speciation node $u$. These variables track the weights of taxon pairs—such as within-branch pairs $w(PA_l^{(g,u)})$ and cross-branch pairs $w(PA_{l,m}^{(g,u)})$—specific to the local topology of node $u$. When a taxon is transferred between partitions, the algorithm avoids a costly global recalculation of $O(D \cdot d)$ complexity. Instead, it identifies the specific nodes affected by the transfer and updates their localized pair weights incrementally in $O(1)$ time.

**Real Taxon Transfer.** A real taxon occupies a single, fixed position in the gene tree topology and contributes a weight of 1. When a real taxon $a$ is transferred from partition $A$ to $B$, the algorithm visits each node $u$ where $a$ is present. If $a$ is located in child branch $i$ relative to node $u$, the localized pair weights at $u$ are updated by removing $a$'s pairings from partition $A$ and establishing its new pairings in partition $B$.

For the within-branch pairs of branch $i$ at node $u$, the updates are:

$$w(PA_i^{(g,u)})_{\text{new}} = w(PA_i^{(g,u)})_{\text{old}} - (w(F_A^{(g,u,i)}) - 1) \tag{2}$$

$$w(PB_i^{(g,u)})_{\text{new}} = w(PB_i^{(g,u)})_{\text{old}} + w(F_B^{(g,u,i)}) \tag{3}$$

For the cross-branch pairs at node $u$, such as those between child branch $i$ and child branch $j$, the updates are:

$$w(PA_{i,j}^{(g,u)})_{\text{new}} = w(PA_{i,j}^{(g,u)})_{\text{old}} - w(F_A^{(g,u,j)}) \tag{4}$$
$$w(PB_{i,j}^{(g,u)})_{\text{new}} = w(PB_{i,j}^{(g,u)})_{\text{old}} + w(F_B^{(g,u,j)}) \tag{5}$$

Analogous $O(1)$ updates are applied at node $u$ to the cross-branch pairs between branch $i$ and the parent branch $k$.

**Dummy Taxon Transfer.** Unlike a real taxon, a dummy taxon $X$ represents a set of taxa from previous subproblems. Relative to a specific speciation node $u$, the constituent taxa of $X$ may be distributed across multiple branches. When $X$ is transferred from partition $A$ to $B$, the algorithm updates every speciation node $u$ where $X$ carries a non-zero weight.

At a given node $u$, let $w_i, w_j$, and $w_k$ be the localized weights of $X$ in branches $i$, $j$, and $k$, respectively. Because $X$ transfers globally between the subproblem partitions, its weights in all branches of node $u$ move simultaneously.

The per-node update for the within-branch pairs in branch $i$ removes $X$'s internal combinations in $A$ and establishes its new combinations in $B$:

$$w(PA_i^{(g,u)})_{\text{new}} = w(PA_i^{(g,u)})_{\text{old}} - w_i \cdot (w(F_A^{(g,u,i)}) - w_i) \tag{6}$$
$$w(PB_i^{(g,u)})_{\text{new}} = w(PB_i^{(g,u)})_{\text{old}} + w_i \cdot w(F_B^{(g,u,i)}) \tag{7}$$

For cross-branch pairs at node $u$ between the two child branches $i$ and $j$, the transfer accounts for $X$'s distributed weights in both branches interacting with the remaining total taxa in the respective partitions:

$$\begin{aligned} w(PA_{i,j}^{(g,u)})_{\text{new}} = w(PA_{i,j}^{(g,u)})_{\text{old}} - \Big(&w_i \cdot (w(F_A^{(g,u,j)}) - w_j) \\ &+ w_j \cdot (w(F_A^{(g,u,i)}) - w_i)\Big) \end{aligned} \tag{8}$$

$$w(PB_{i,j}^{(g,u)})_{\text{new}} = w(PB_{i,j}^{(g,u)})_{\text{old}} + w_i \cdot w(F_B^{(g,u,j)}) + w_j \cdot w(F_B^{(g,u,i)}) \tag{9}$$

By performing these updates locally at each affected node $u$, the algorithm efficiently handles simultaneous weight changes across multiple branches. This per-node approach ensures the FM heuristic remains fast and scalable, as it avoids the need to recalculate scores from scratch for the entire subproblem.

#### 2.2 Normalization in wQFM-TREE and wQFM-GDL-T

##### 2.2.1 Normalization in wQFM-TREE

We discuss the normalization scheme used in TREE-QMC and wQFM-TREE in detail and explain how it is applied in wQFM-TREE using illustrative examples. As explained in the main text, the need for normalization arises from the introduction of dummy taxa within subproblems during the divide phase. Let's consider Subproblem B in Figure 2. It contains taxa $1, 2, 4, 6, 7, 8, 9, X$, but the gene trees don't contain $X$. From the perspective of this subproblem, we realize that $X$ represents 3 and 5, the taxa of the sister subproblem. Thus, for algorithmic calculations of this subproblem, the leaves 3 and 5 in the gene trees are relabeled with $X$. However, as multiple taxa are represented by $X$, this will inflate the count of quartets associated with $X$. For example, quartets $1, 2|4, 3$ and $1, 2|4, 5$ both will be considered as $1, 2|4, X$. Thus, when scoring candidate bipartitions for Subproblem B using satisfied and violated quartets, quartets containing $X$ will have a higher weight than quartets containing only real taxa, negatively affecting the results. To mitigate this effect, we have to downweight these quartets.

To implement the normalization scheme, each taxon is assigned a weight, and the weight of a quartet $ab|cd$ is defined as the product of the weights of its four taxa, $w(ab|cd) = w(a).w(b).w(c).w(d)$. The real taxa are assigned weight 1, and thus the quartets containing real taxa also receive unit weight. However, to downweight the quartets associated with dummy taxa, the real taxa represented by a dummy taxon are each assigned a weight less than 1, and the sum of their weights equals 1. In this scenario, taxa 3 and 5 are each assigned a weight of 1/2. As a result, the quartets $1, 2|4, 3$ and $1, 2|4, 5$ each receive a weight of 1/2, and together they sum to 1, thereby mitigating the effect.

Now, in a more complex scenario such as in Subproblem C, a dummy taxon $Y$ in $C$ represents $1, 7$, and $X$ and $X$ represents 3 and 5 as before. Thus, taxa 3 and 5 are getting relabelled multiple times for this subproblem. Now, for dummy taxon $Z$, the normalization works the same as for dummy taxon $X$ in subproblem B. However, for $Y$, an important point to note is that the real taxa under $Y$, taxa $1, 3, 5, 7$ are not weighted uniformly (Figure 2). 1 and 7 are directly represented by $Y$ similarly as $X$. Thus, the unit weight allocated for $Y$ in that subproblem is distributed equally among them. Consequently, $X$ receives weight 1/3, which is equally distributed to 3 and 5. Thus, the taxa are weighted non-uniformly depending on the subproblem decomposition in the recursion tree, and taxa that are more relevant to a particular subproblem are assigned higher weights.

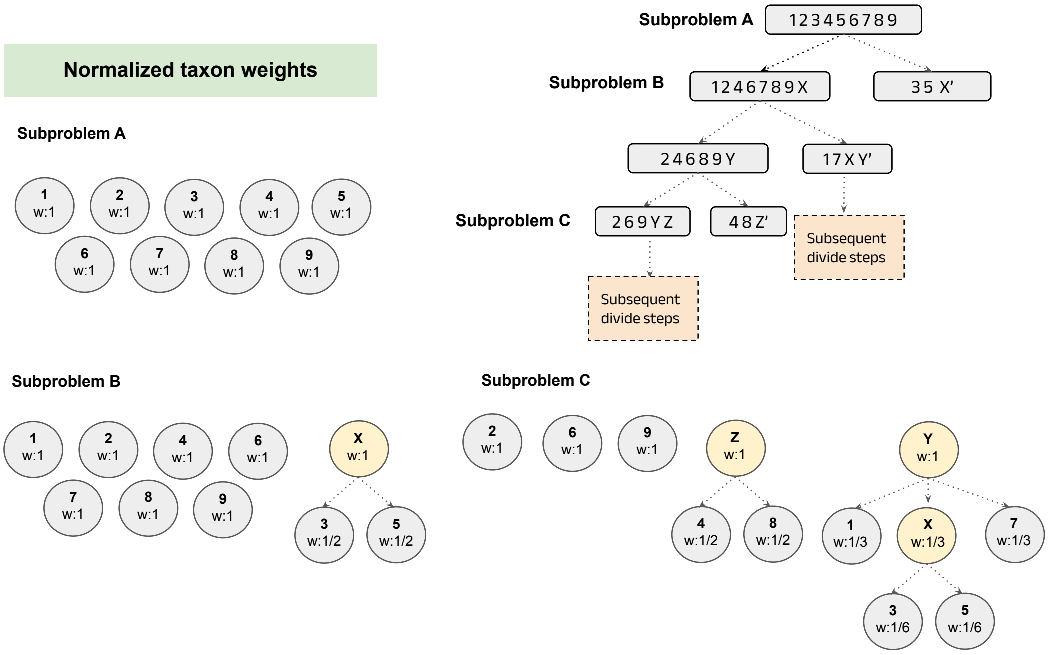

Figure 2: **Normalized taxon weights for different subproblems in the example divide phase of wQFM-TREE.** Real and dummy taxa in each subproblem are assigned a weight of 1, and the weights of the taxa represented by a dummy taxon are normalized so that their total sums to 1. For subproblem A, no dummy taxon is present, and normalization has no effect. For subproblem B, this is a simple case where the dummy taxon $X$ represents taxa 3 and 5. For subproblem C, we observe that the taxa represented by dummy taxon $Y$ are not assigned equal weights. Instead, they are weighted based on their relevance to the subproblem and subproblem structure. Since taxa 3 and 5 under dummy taxon $Y$ are not directly represented by $Y$, but rather through $X$, they receive lower weights compared to taxa 1 and 7.

##### 2.2.2 Normalization in wQFM-GDL-T

The normalization scheme of wQFM-TREE can be directly applied to wQFM-GDL, but we enhance it with a locus-aware scheme. The main philosophy is that we do not apply the same normalization factors across the entire gene family tree. Instead, we compute separate normalization factors for different locus-specific subtrees—an approach we call the locus-aware normalization scheme (Figure 3). This is motivated by the fact that each locus undergoes a distinct history of duplication and loss, and the set of taxa present in each locus-specific subtree can vary. Since normalization factors depend on the taxa set, they naturally differ across loci. Consequently, quartets defined on the same four taxa $a, b, c, d$ may receive different weights depending on the locus from which they originate.

The key idea in the implementation is that all quartets on $a, b, c, d$ forming SQ equivalent class descend from the same ancestral locus at the time of the speciation event corresponding to their anchor LCA (1) and for each group of quartet equivalence classes associated with a particular anchor LCA (which reside in the locus-specific subtrees associated with that locus), we compute normalization factors separately and independently. To determine the taxa set relevant for these quartets, we identify the branches and subtrees that can contribute to forming them. For a speciation node acting as the anchor LCA, the taxa set includes (i) the taxa in the subtrees rooted at that node and (ii) the taxa associated with branches of the speciation nodes along the path from the anchor LCA to the root. Taxa outside these regions cannot participate in forming the corresponding quartets and are excluded. The resulting restricted taxa set is then used to compute normalization factors specific to that group of quartets.

It is important to note that this is an approximate strategy rather than an exact formulation. The goal is to reduce the bias introduced by the traditional normalization scheme while keeping the computation tractable. Although more sophisticated approaches could be designed, our experimental observations suggest that they would likely increase time complexity without providing meaningful gains in accuracy.

In Figure 3, duplication in the $dup1$ node creates a new locus, and both evolve in parallel and independently. In case of one of the loci that evolves along the left subtrees, $f$ taxon is absent due to a loss event. All quartets with anchor LCA in the blue region are restricted to taxa set $a, b, c, d, e, g, h$ as there is no $f$ taxon, which results in the normalized taxon weights as shown in the Figure. On the other hand, in the right subtree, $c, d$ is absent. Moreover, there is another duplication event at the node $dup2$. For quartets with anchor LCA in that right subtree, the effective taxa set is $a, b, e, f, g, h$. Thus, the normalization factors should be changed.

#### 2.3 wQFM-GDL-Q: Details and Pseudocode

Following an approach similar to the score calculation process of a candidate bipartition in wQFM-GDL-T, for each speciation node in the tagged and rooted gene trees, we independently generate the speciation-driven quartets with anchor LCA at that particular node and add them to the set. Similar to wQFM, the weight of a quartet is defined as its frequency across the gene trees. However, because the gene trees here are multicopy, multiple instances of the same quartet may occur within a single gene tree, and consequently, the total frequency (weight) of a quartet can be substantially larger than the number of gene trees, unlike in wQFM.

Now, we consider enumerating all SQs with a particular anchor LCA $u$ with child branches $i$ and $j$. For balanced quartets, we enumerate all possible taxon pairs formed within each child branch. Of course, we keep only one copy of a taxa pair as they form equivalent quartets, and we must consider one representative from them. Then, any pair from branch $i$ can create a valid quartet with any pair from branch $j$. We generate all

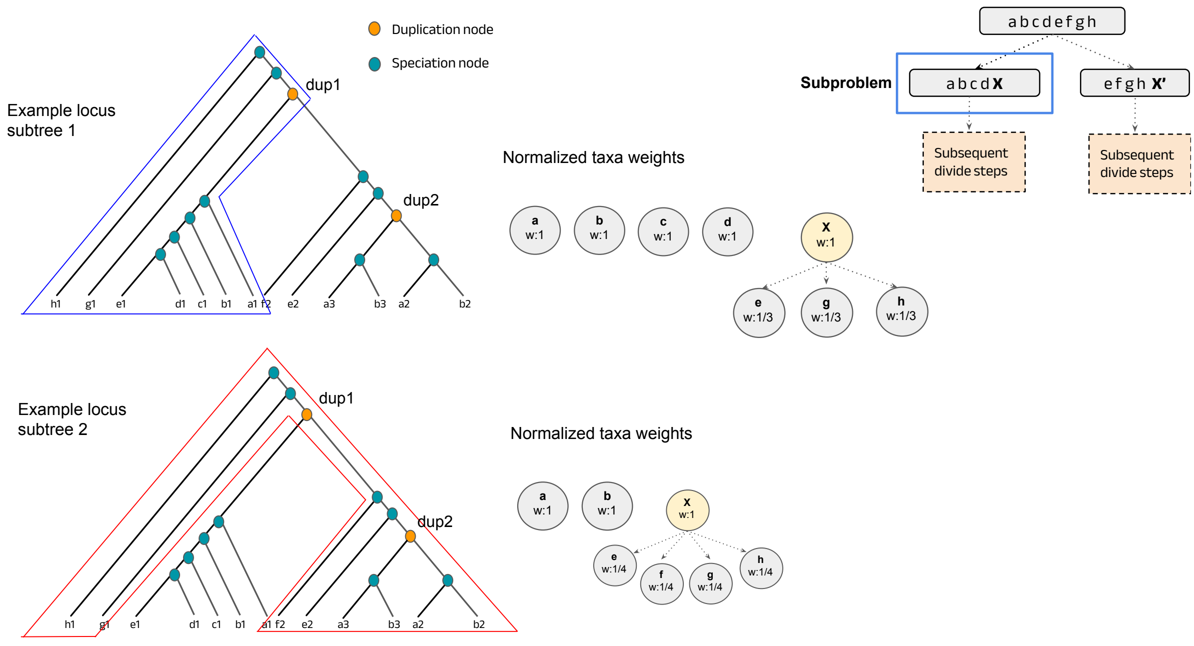

Figure 3: **Different normalized taxon weights for quartets associated with different regions of the gene family tree.** All quartets with anchor LCA in the blue region are restricted to taxa set $a, b, c, d, e, g, h$ as $f$ taxon is absent due to a loss event. Similarly, quartets with anchor LCA in the red region are restricted to the taxa set $a, b, e, f, g, h$. Thus, the normalization factors vary within the gene family tree.

possible quartets by joining the pairs.

For the unbalanced quartets, one pair of taxa will be present in one of the child branches $i$ or $j$. We generate all possible pairs for both child branches. For the second pair, one taxon will be present in the other child branch and one taxon in the parent branch. As discussed, not all taxa in the parent branch are considered here (Figure 2 in the main paper). We list all taxa present in the child branches and the parent branch and merge them to create all valid second pairs. The pairs are then merged appropriately to enumerate quartets.

An important point to note is that, although the high-level idea resembles the score calculation procedure of wQFM-GDL-T, there is no notion of satisfied or violated quartets here because there is no bipartition to consider at this stage. Instead, our objective is to generate all speciation-driven quartets (SQs) from the gene trees so that wQFM can operate on them. Since this step focuses on explicit quartet enumeration rather than efficiently computing scores directly from the gene trees, the procedure is more straightforward compared to wQFM-GDL-T. The algorithm for generating the quartet set is shown in Algorithm 1.

**Algorithm 1:** Algorithm for generating the quartet set for wQFM-GDL-Q.

**Input :** $\mathcal{G}$, set of tagged and rooted multicopy gene family trees

**Output:** The set of speciation-driven quartets in $\mathcal{G}$ with only one representative for each equivalence class

```
Q ← ∅ ;                                          // quartet set initialization
foreach gene tree g ∈ G do
    foreach internal speciation node s in g do
        l ← left child of s
        r ← right child of s

        qleft ← getRealTaxainSubtree(l) // list all taxa present in the
            subtree
        qright ← getRealTaxainSubtree(r)
        pairs_left ← getAllPairs(qleft) // generate all possible taxa pairs
            from the taxa list
        pairs_right ← getAllPairs(qright)
        Q1 ← generateQuartetsfromPairs(pairs_left,pairs_right)
        /* generate all valid quartets (pair of pairs) from the two lists of
            pairs                                                         */

        par ← getRealTaxainParents(s)
        /* list all the taxa present in the other child subtree of all the
            speciation nodes on the path from s to root                   */
        crosspairs_left ← getAllCrossPairs(par,qleft) // generate all
            possible pairs with one taxa from each of the taxa lists
        crosspairs_right ← getAllCrossPairs(par,qright)
        Q2 ← generateQuartetsfromPairs(pairs_left,crosspairs_right)
         ∪generateQuartetsfromPairs(crosspairs_left,pairs_right)

        Q ← Q ∪ Q1 ∪ Q2
    end
end
return Q
```

```
Function getRealTaxainParents(s)
    taxa_list ← ∅
    current ← s
    while current.parent ≠ null do
        if current.parent is speciation node then
            otherChild ← getSibling(current)
            taxa_list ← taxa_list ∪ getRealTaxainSubtree(otherChild)
        end
        current ← current.parent
    end
    return taxa_list
```

### 3 Simulation details and parameters

We simulated two large datasets, SIM200 and SIM500, containing model conditions of 200 and 500 taxa, respectively. In this section, we describe the simulation procedure in detail, including the commands used and the parameter settings. For both datasets, we varied three duplication rates. For each duplication rate, we considered two loss rates, two levels of ILS, and different numbers of gene trees (250, 500, and 1000). This resulted in a total of 36 model conditions per dataset, with 20 replicates per model condition for SIM200 and 10 replicates for SIM500. Simulation settings are summarized in Table 1.

#### 3.1 True gene family tree simulation

We used Simphy (2), which is a widely used program for simulation of gene family evolution under incomplete lineage sorting (ILS), gene duplication and loss (GDL), and horizontal gene transfer (HGT). The simulation parameters used in Simphy are presented in detail in Table 2.

For both SIM200 and SIM500, we varied the duplication rate to create model conditions with mean numbers of extra gene copies per species of 0.25, 1, and 3. That means, due to duplication and loss, the average numbers of gene copies per species are 1.25, 2, and 4, respectively. In SIM500, the model condition with an average of 3 extra copies per species contains gene trees with significant duplication, having more than 2,000 leaves on average. To determine the appropriate duplication rates in Simphy for generating these conditions, we systematically varied the duplication rate and, for each value, generated 1000 replicates for datasets with 200 taxa (SIM200) and 500 taxa (SIM500), using a loss rate of zero. As expected, the mean number of copies increased with higher duplication rates. We then identified the precise duplication rate that produced the desired mean copy number (0.25, 1, or 3) using a binary-search style procedure over the duplication parameter space. The loss rate is varied for each duplication rate simply as the ratio of

Table 1: Simulation settings for SIM200 and SIM500 with varying parameters. For each of the duplication rates, we vary the loss rate, ILS level, and number of gene trees.

| **Condition** | **Parameter Ranges** |
|---|---|
| No. of taxa | 200,500 |
| Varying no. of gene trees | 250, 500, and 1000 |
| Varying duplication rate | 0.25, 1, and 3 (Mean number of extra copies per species) Corresponding duplication rates for S200: $\{0.8, 2.0, 4.1\} \times 10^{-10}$ events/generation Corresponding duplication rates for S500: $\{0.8, 2.1, 4.3\} \times 10^{-10}$ events/generation |
| Varying loss rate | 0.1 and 1 (fraction of duplication rates) |
| Varying ILS | 25% and 70% (mean RF distance between true gene trees and the species tree ) Corresponding haploid effective population size for S200: $\{0.5, 1.3, 2.9\} \times 10^8$ Corresponding haploid effective population size for S500: $\{0.5, 1.25, 2.7\} \times 10^8$ |
| Sequence length | 100bp |

0.1 to 1 of the duplication rate. In the case of a 0.1 ratio, there is a small number of loss events, and the number of copies is high. In a ratio of 1, there is a significant number of loss events, and the mean number of copies is closer to the number of taxa, even with high duplication rates.

Similarly, for each duplication rate, we simulated two conditions varying ILS: 25% and 70% ILS, meaning the average RF rates of true gene trees and species tree are 25% and 70%, respectively. As the amount of ILS increases, the RF rate goes higher. We estimated the proper haploid effective population sizes for these two conditions in a procedure similar to the estimation of duplication rates.

**Simphy command for SIM200 with standard parameters:**

```
simphy -sl f:200 -rs 20 -rl f:1000 -rg 1 -sb f:0.000000005 -sd f
   :0 -st ln:21.25,0.2 -so f:1 -si f:1 -sp f:435000000 -su ln:
   21.9,0.1 -hh f:1 -hs ln:1.5,1 -hl ln: 1.551533, 0.6931472 -hg
   ln:1.5,1 -cs 9644 -v 3 -o default -ot 0 -op 1 -lb f
```

Table 2: Simphy simulation Parameters for SIM200 and SIM500 dataset

| Parameter name | Parameter value |
|---|---|
| **Standard Parameters for SIM200/500** | |
| No. of taxa | 200/500 + outgroup |
| Speciation rate | 5e-9 |
| Extinction rate | 0 |
| Locus trees | 1000 |
| Gene trees | 1 |
| No. of replicates | 20/10 |
| Ingroup divergence to the ingroup ratio | 1.0 |
| Generations | LogN(21.25, 0.2) |
| Haploid effective population size | 2.9e+8/2.7e+8 |
| Global substitution rate | LogN(-21.9, 0.1) |
| Lineage specific rate gamma shape | LogN(1.5, 1) |
| Gene family specific rate gamma shape | LogN(1.551533, 0.6931472) |
| Gene tree branch specific rate gamma shape | LogN(1.5, 1) |
| Duplication rate | 4.1e-10/4.35e-10 |
| Loss rate to duplication rate ratio | 1 |
| Seed | 9644 |
| **Varying Duplication and Loss Rates - SIM200** | |
| Duplication rate | 0.8e-10, 2e-10, 4.1e-10 |
| Loss rate to duplication rate ratio | 0.1, 1 |
| **Varying Duplication and Loss Rates - SIM500** | |
| Duplication rate | 0.8e-10, 2.1e-10, 4.35e-10 |
| Loss rate to duplication rate ratio | 0.1, 1 |
| **Controlling Duplication and ILS Rate - SIM200** | |
| Duplication rate | 0.8e-10, 2e-10, 4.1e-10 |
| Haploid effective population size | 0.5e+8, 2.9e+8 |
| **Controlling Duplication and ILS Rate - SIM500** | |
| Duplication rate | 0.8e-10, 2e-10, 4.35e-10 |
| Haploid effective population size | 0.5e+8, 2.7e+8 |

```
:0.00000000041 -ld f:0.00000000041 -lt f:0
```

#### 3.2 Simulating MSA

We use Alisim (3) to generate sequence alignments from the simulated true trees. The parameters are presented in Table 3.

Table 3: Simulation Parameters for AliSim

| **Parameter name** | **Parameter value** |
|---|---|
| Sequence length | 100 |
| Sequence base frequencies | Dirichlet(A=36, C=26, G=28, T=32) |
| Sequence transition rates | Dirichlet(TC=16, TA=3, TG=5, CA=5, CG=6, AG=15) |
| Seed | 9644 |

AliSim is implemented in IQ-TREE version 2.4.0 or later. We use IQ-TREE to generate the sequence alignments.

**IQ-TREE command for SIM200 with standard parameters:**

```
iqtree3 --alisim SimPhy_1.0.2/bin/default/1/MSA1 -t tree_file -m
   "GTR{1/3/0.6/1.2/3.2}+F
   {0.2950819672/0.2131147541/0.2295081967/0.2622950820}" --
   seqtype DNA --length 100 --seed 9644
```

#### 3.3 Simulating estimated gene trees from MSA

We use FastTree (4) to estimate maximum-likelihood gene trees from the sequence alignment under the general time-reversible model of nucleotide substitution with four discrete gamma rates (GTR+G4). We used the simple command:

```
FastTree -gtr -gamma -nt alignment_file > tree_file
```

### 4 Running time and memory consumption

Table 4: Running time (in minutes) of ASTRAL-Pro3, DupLoss-2, SpeciesRax, FastMulRFS and wQFM-GDL-T under a few model conditions with low, moderate, and high duplication and loss across different numbers of gene trees (SpeciesRax could not complete all model conditions within our computational budget). All methods were executed on 5 replicates for each model condition to measure running time, and the average, along with the standard deviation, is shown for each model condition.

| **Model condition** | **Method** | **Number of gene trees** | | |
|---|---|---|---|---|
| | | **250** | **500** | **1000** |
| **sim200_dup1_loss1** | ASTRAL-Pro3 | 2.86 ± 0.36 | 6.80 ± 0.73 | 9.73 ± 0.49 |
| | DupLoss-2 | 1.49 ± 0.09 | 3.73 ± 0.25 | 9.85 ± 1.13 |
| | SpeciesRax | 279.27 ± 22.10 | – | – |
| | FastMulRFS | 1.95 ± 0.39 | 6.50 ± 1.44 | 20.39 ± 6.63 |
| | wQFM-GDL-T | 4.58 ± 0.75 | 11.92 ± 1.17 | 23.27 ± 1.98 |
| **sim200_dup0_ILS25** | ASTRAL-Pro3 | 2.73 ± 0.25 | 6.40 ± 0.43 | 11.00 ± 0.82 |
| | DupLoss-2 | 0.51 ± 0.01 | 1.36 ± 0.02 | 3.41 ± 0.10 |
| | SpeciesRax | 163.59 ± 15.69 | – | – |
| | FastMulRFS | 0.71 ± 0.20 | 1.61 ± 0.64 | 3.58 ± 1.56 |
| | wQFM-GDL-T | 0.85 ± 0.23 | 2.36 ± 0.33 | 4.39 ± 0.53 |
| **sim200_dup3_loss0.1** | ASTRAL-Pro3 | 4.30 ± 0.76 | 11.00 ± 1.55 | 26.40 ± 3.31 |
| | DupLoss-2 | 2.50 ± 0.13 | 6.26 ± 0.38 | 16.92 ± 1.49 |
| | SpeciesRax | 329.18 ± 20.58 | – | – |
| | FastMulRFS | 1.11 ± 0.19 | 2.76 ± 0.48 | 7.91 ± 1.68 |
| | wQFM-GDL-T | 13.15 ± 1.69 | 34.24 ± 2.84 | 68.44 ± 4.15 |
| **sim500_dup1_loss1** | ASTRAL-Pro3 | 14.76 ± 1.53 | 31.87 ± 2.84 | 55.73 ± 4.97 |
| | DupLoss-2 | 20.28 ± 0.67 | 45.89 ± 1.87 | 127.15 ± 8.85 |
| | SpeciesRax | – | – | – |
| | FastMulRFS | 13.66 ± 1.70 | 39.45 ± 5.84 | 124.22 ± 46.14 |
| | wQFM-GDL-T | 118.04 ± 16.29 | 277.73 ± 25.43 | 568.88 ± 45.12 |
| **sim500_dup0_ILS25** | ASTRAL-Pro3 | 11.87 ± 0.55 | 31.13 ± 1.04 | 55.87 ± 1.73 |
| | DupLoss-2 | 8.31 ± 0.31 | 26.35 ± 0.76 | 67.93 ± 4.13 |
| | SpeciesRax | – | – | – |
| | FastMulRFS | 5.33 ± 3.04 | 13.15 ± 7.63 | 26.81 ± 5.40 |
| | wQFM-GDL-T | 3.94 ± 1.25 | 12.98 ± 1.72 | 24.75 ± 3.51 |
| **sim500_dup3_loss0.1** | ASTRAL-Pro3 | 33.95 ± 10.72 | 89.00 ± 22.48 | 234.25 ± 63.97 |
| | DupLoss-2 | 43.63 ± 0.89 | 95.17 ± 2.60 | 279.63 ± 12.94 |
| | SpeciesRax | – | – | – |
| | FastMulRFS | 6.60 ± 1.44 | 14.65 ± 5.36 | 33.21 ± 5.96 |
| | wQFM-GDL-T | 264.17 ± 26.20 | 599.06 ± 43.28 | 1301.28 ± 80.55 |

The running times of all methods on the large datasets are reported in Table 4. All experiments were conducted using a single core of an AMD Ryzen 9 7950X processor and 64 GB RAM. In our machine, ASTRAL-Pro3 or FastMulRFS seems to be the fastest method on most model conditions, while SpeciesRax is the slowest. A point to note is that, ASTRAL-Pro3 and SpeciesRax can achieve good parallel efficiency and may run faster in environments that better support full parallel processing, which was not available in our setup. The running time of wQFM-GDL is comparable to ASTRAL-Pro3 and DupLoss-2, though it is typically slower. However, it successfully analyzed 500-taxon model conditions with a high duplication rate (more than 2,000 leaves per gene family tree) in 20 hours, showing that it is scalable enough to handle large datasets. The QFM framework is inherently parallelizable, and future works should focus on CPU and GPU parallelization of wQFM-GDL to improve its runtime performance.

wQFM-GDL successfully analyzed the large 200-taxon and 500-taxon datasets using no more than 8 GB and 16 GB of memory, respectively. FastMulRFS required substantially higher memory, using around 22 GB, while ASTRAL-Pro3 and DupLoss-2 were more memory-efficient, requiring approximately 2 GB and 800 MB of memory, respectively, for both datasets.

### 5 Additional results

#### 5.1 Running commands for the methods

**Command for wQFM-GDL-T:**

```
java -jar wqfm-gdl.jar -i INPUT_FILE -o OUTPUT_FILE -t
```

**Command for wQFM-GDL-Q:**

```
java -jar wqfm-gdl.jar -i INPUT_FILE -o OUTPUT_FILE -q
```

**Command for ASTRAL-Pro3:**

```
astral-pro3 -o OUTPUT_FILE INPUT_FILE 2>LOG_FILE
```

**Command for SpeciesRax:**

```
mpiexec -np 1 generax --families families.txt --species-tree
   MiniNJ --strategy SKIP --rec-model UndatedDTL --per-family-
   rates --prune-species-tree --si-strategy HYBRID
```

**Command for DupLoss-2:**

```
./DupLoss-2.out -i INPUT_FILE -o OUTPUT_FILE
```

**Command for FastMulRFS:**

```
python ../python-tools/preprocess_multrees_v3.py \
    -i g_trees-mult.trees \
    -o g_trees-mult-for-fastrfs.trees \
    --verbose
../external/FastRFS/build/FastRFS \
        -i g_trees-mult-for-fastrfs.trees \
        -o fastmulrfs.tree &> fastmulrfs.log
```

To run FastMulRFS, we used the scripts available at the FastMulRFS' Github repository (https://github.com/ekmolloy/fastmulrfs/tree/master/example).

The results of SpeciesRax and DupLoss-2 may vary depending on the random seed. Both methods were run once on each input using their default settings.

#### 5.2 Results Including DupLoss-2 and FastMulRFS

The results of S25 including DupLoss-2 and FastMulRFS and the results of SIM200 and SIM500 datasets including DupLoss-2 are presented in Figures 4 and 5. These methods are excluded from the corresponding figures of the main paper for clearer visualization, as their error rates are substantially higher than those of the other methods. The results of DupLoss-2 vary across different random seeds. It was run once on each input using default parameter settings. Most model conditions in these datasets include both ILS and GDL. We observe that it performs reasonably well under low ILS conditions, but its accuracy declines substantially as the ILS level increases.

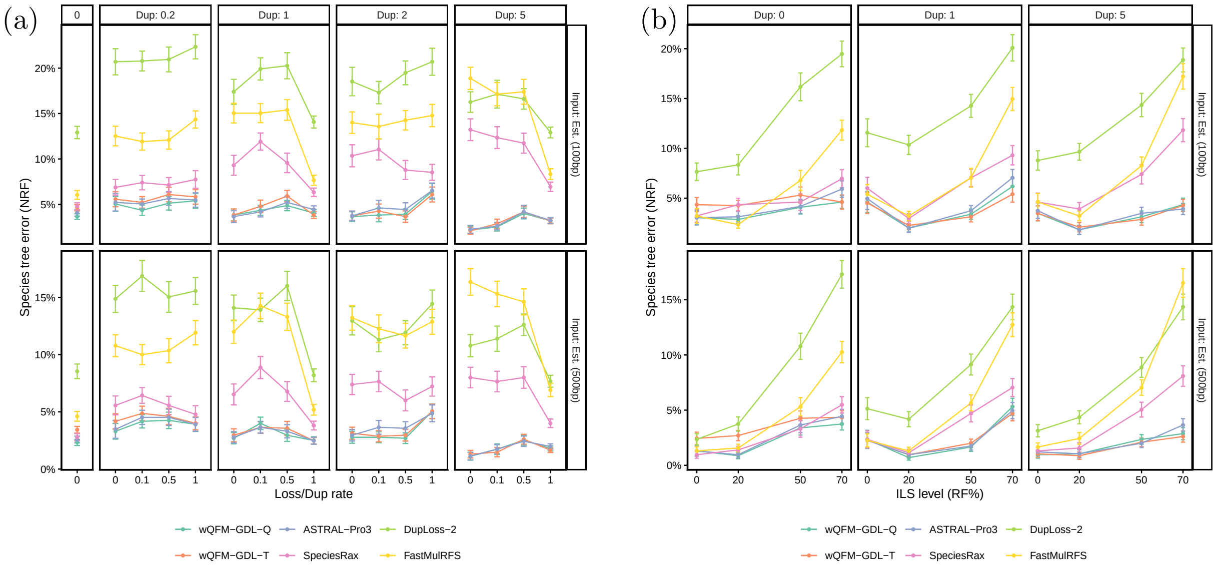

Figure 4: Species tree error on the S25 dataset, including DupLoss-2 and FastMulRFS.(a) Controlling duplication rate (box columns, labelled by mean number of copies per species minus one) and loss rate (labelled by ratio of loss and duplication rate). (b) Controlling the duplication rate and ILS level (RF rate between true gene trees and species tree).

#### 5.3 Empirical Datasets

##### 5.3.1 Vertebrates188 dataset

The full tree and some important subtrees recovered by wQFM-GDL for the Vertebrates188 dataset are shown below.

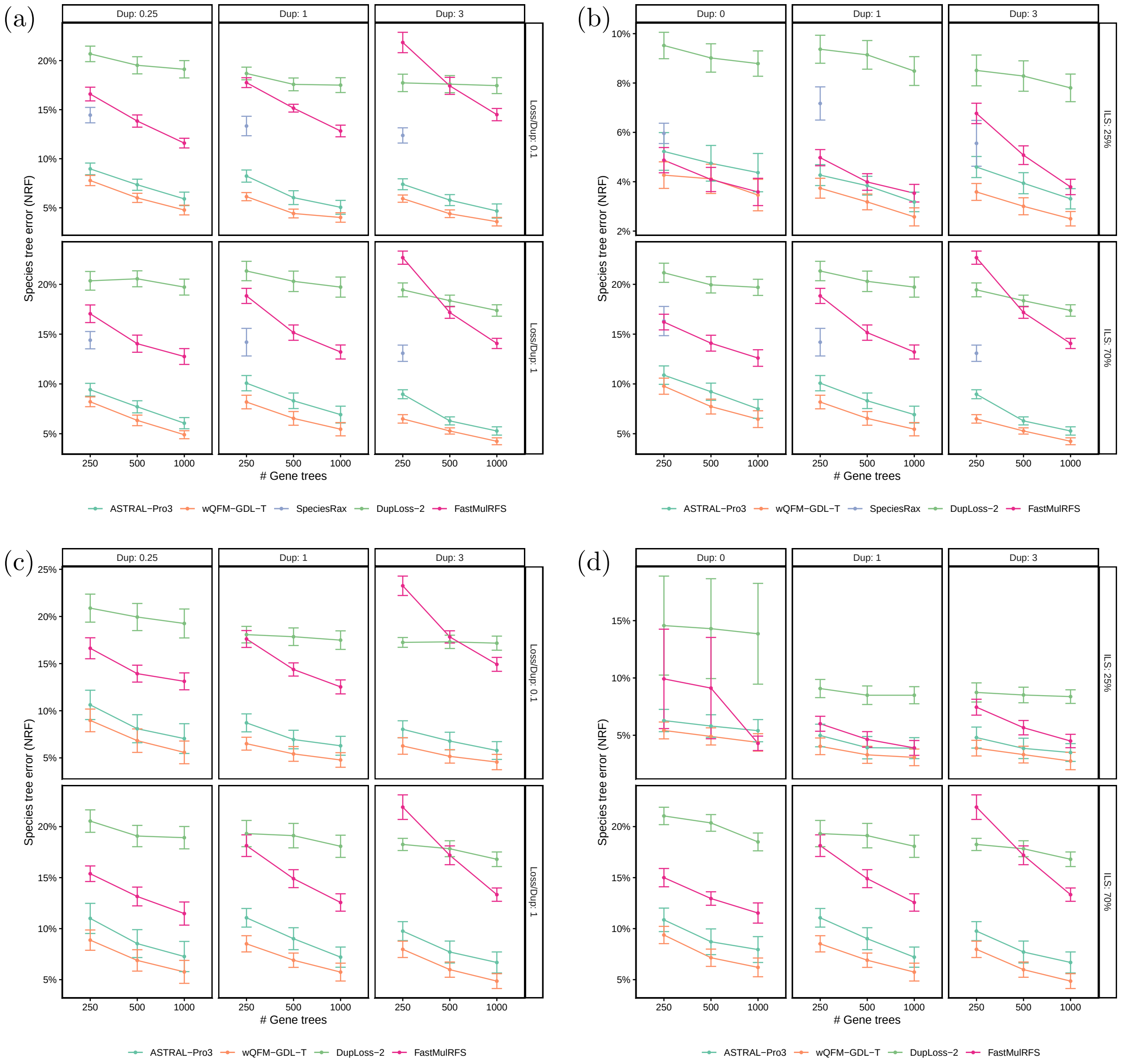

Figure 5: Species tree error on the SIM200 (a-b) and SIM500 (c-d), including DupLoss-2. (a,c) Controlling duplication and loss rate. (b,d) Controlling the duplication rate and ILS level.

##### 5.3.2 Archaea364 dataset

The species trees inferred with wQFM-GDL and ASTRAL-Pro3 from the Archaea364 dataset using the full set, top 50% and 25% marker proteins are shown below.

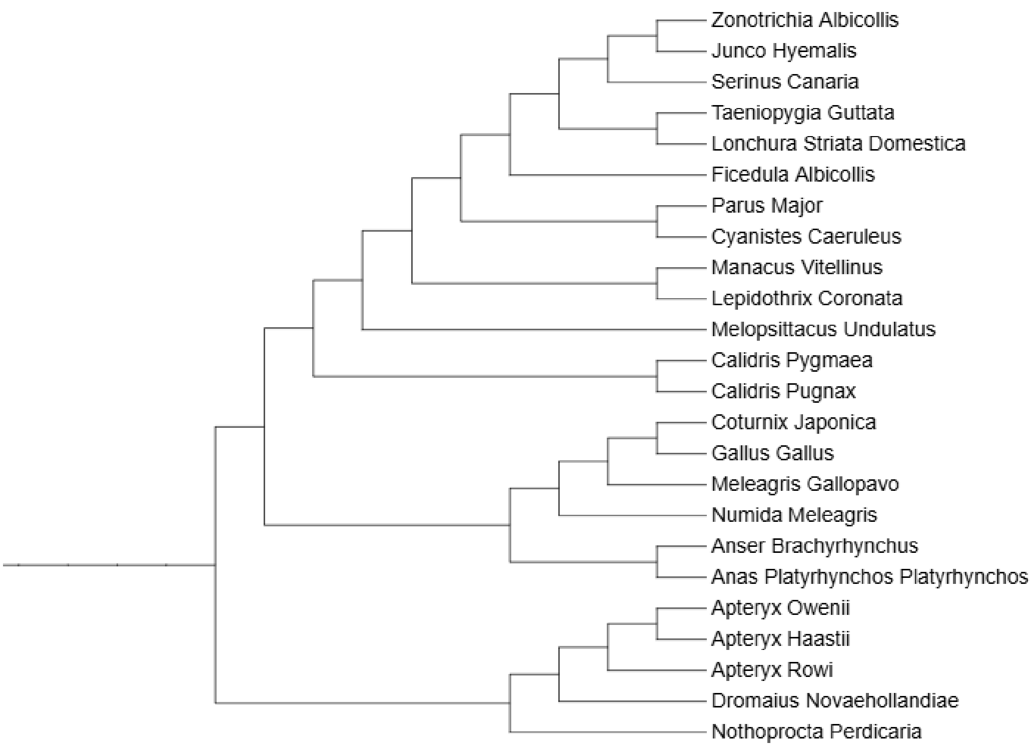

(a) The avian subtree

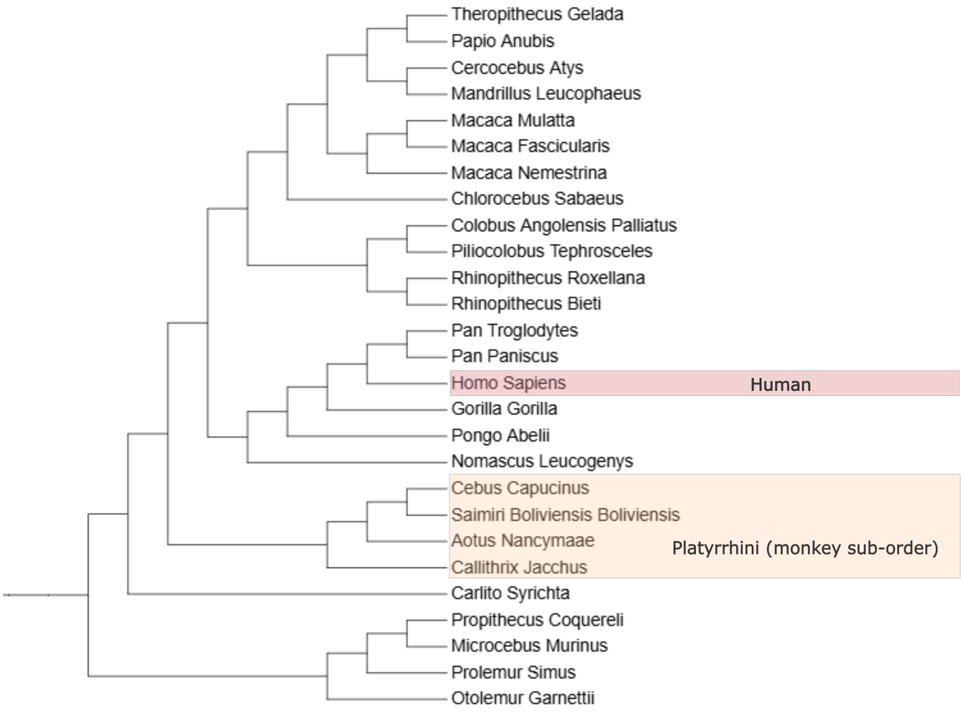

(b) The primates subtree and the contentious monkey sub-order

Figure 6: Important contentious subtrees of the species tree estimated by wQFM-GDL for the Vertebrates188 dataset

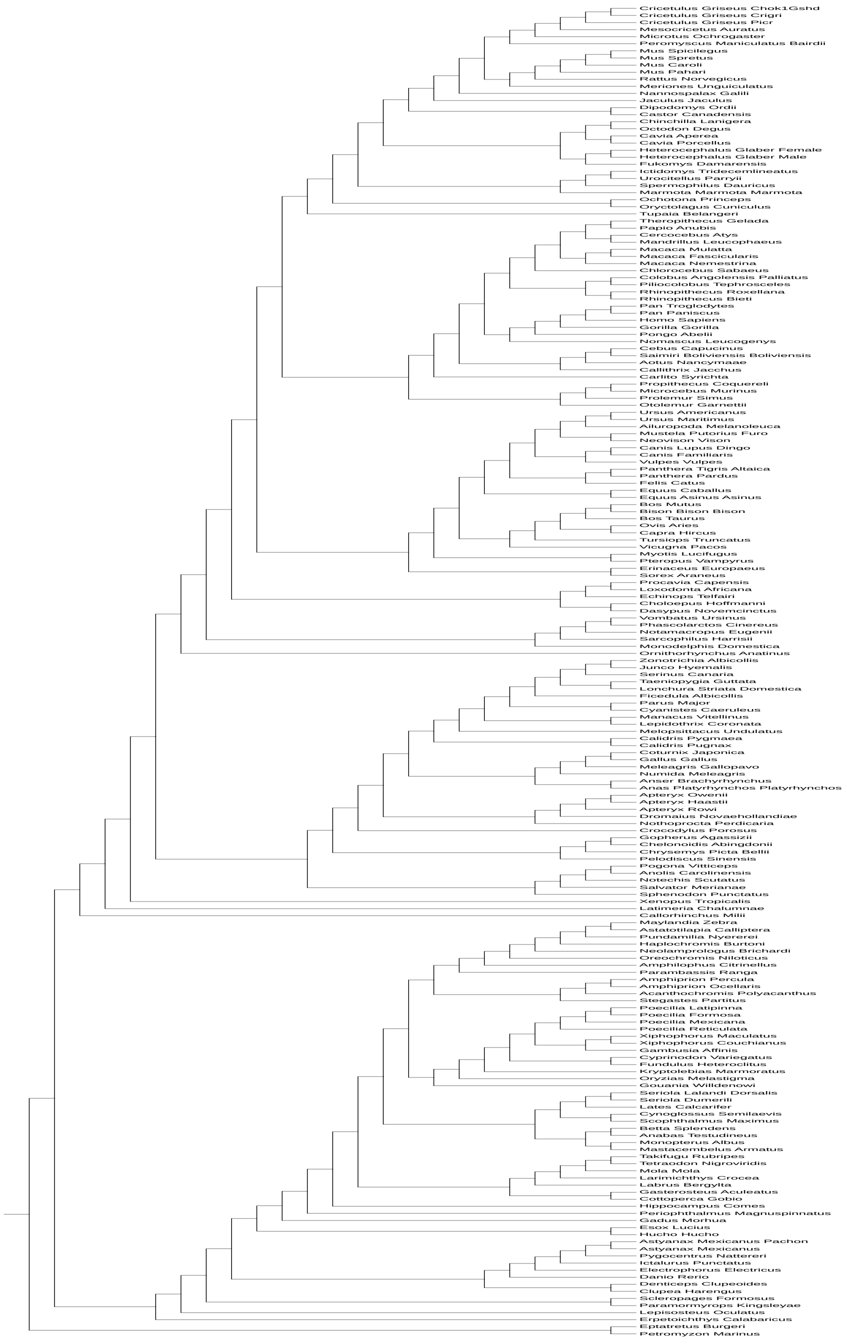

Figure 7: The species tree inferred with wQFM-GDL for the Vertebrates188 dataset.

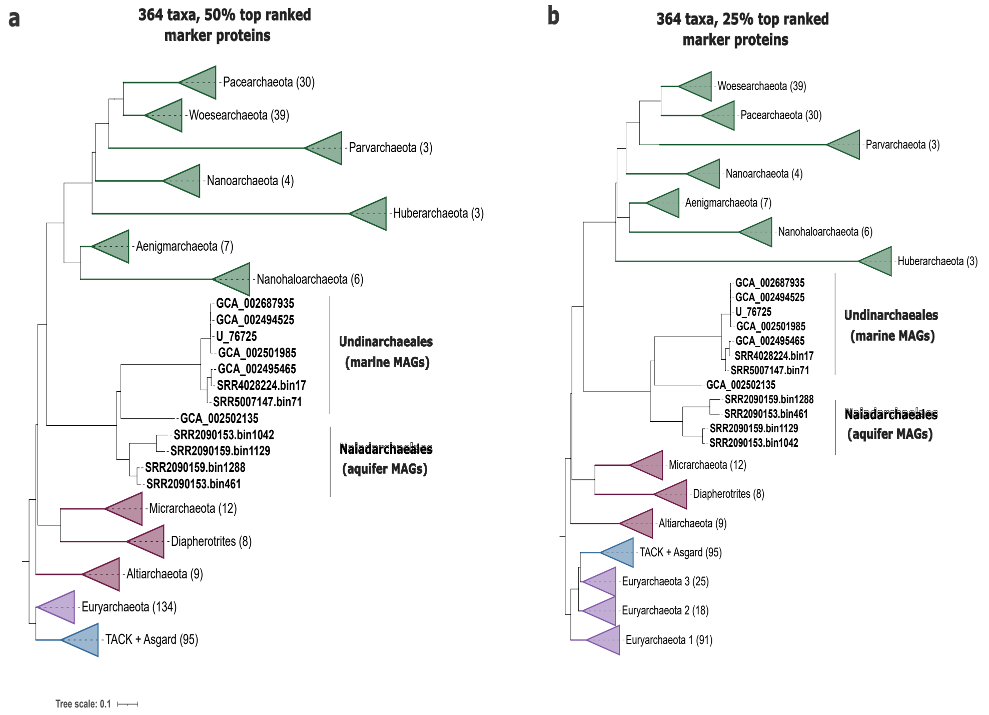

Figure 8: The species trees inferred with wQFM-GDL from the Archaea364 dataset with top 50% and 25% marker proteins

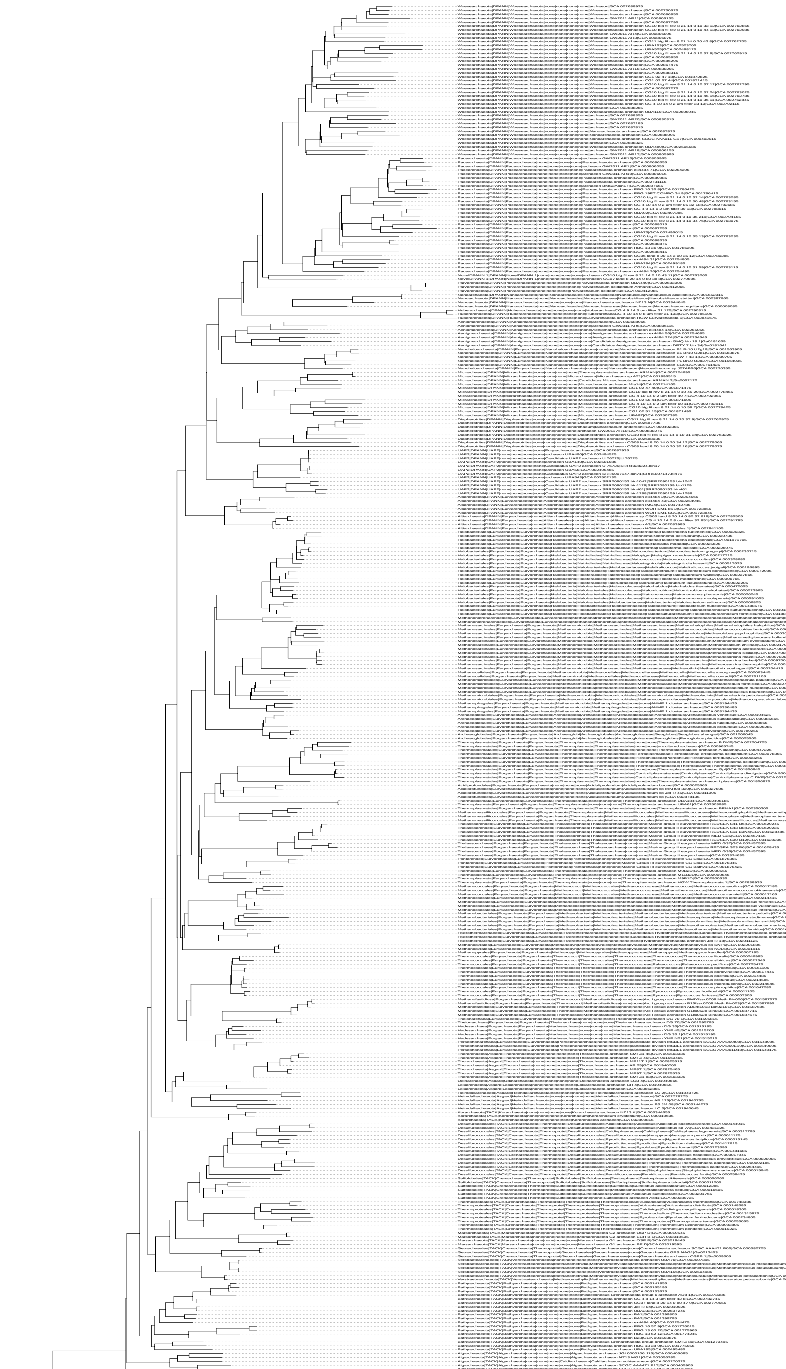

Figure 9: The species tree inferred with wQFM-GDL for the Archaea364 dataset using the full set of protein markers.

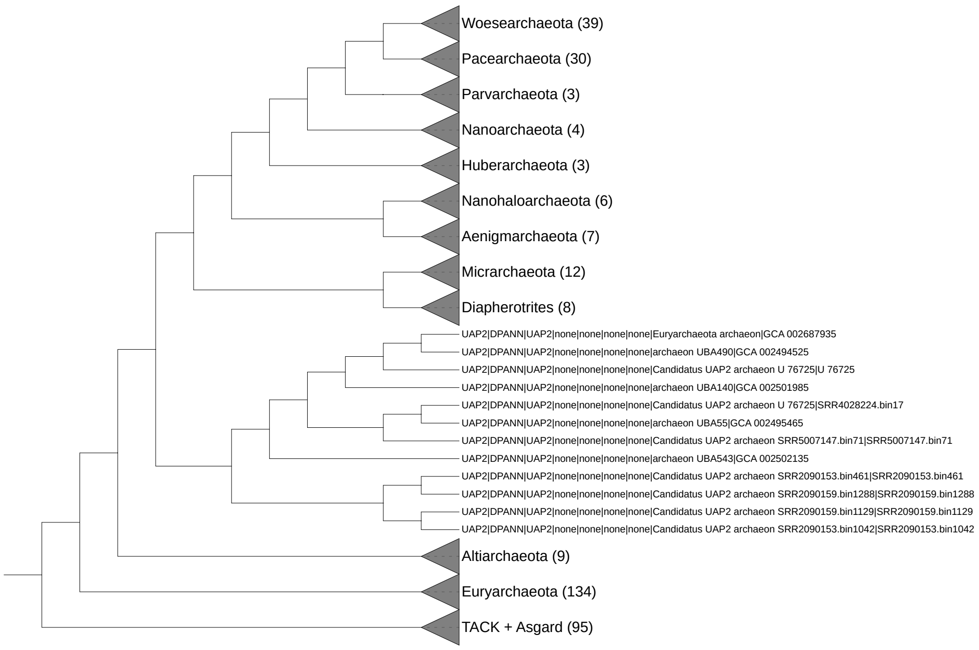

(a) The species tree using the top 50% protein markers.

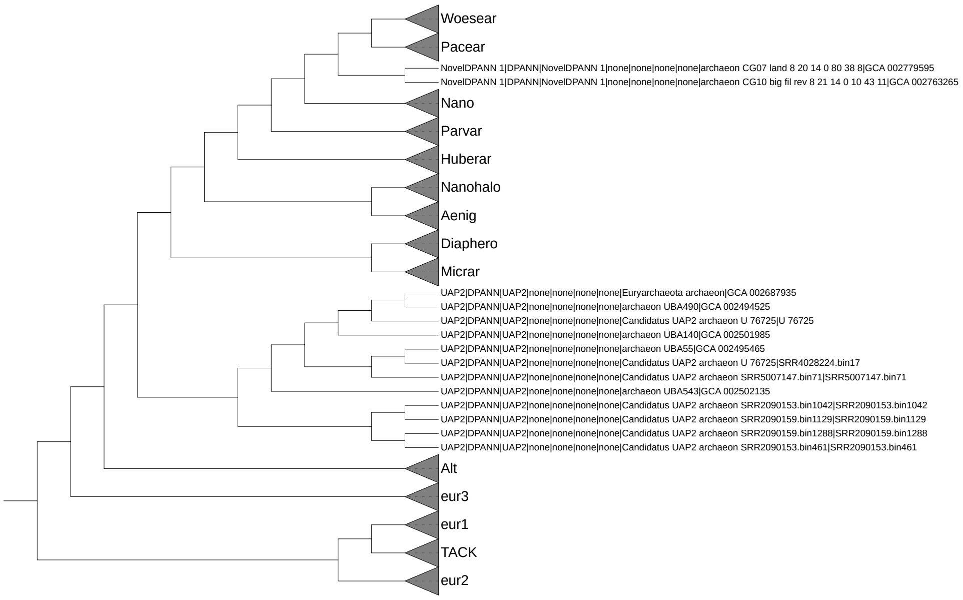

(b) The species tree inferred using the top 25% protein markers.

Figure 10: The species tree inferred with ASTRAL-Pro3 for the Archaea364 dataset
